# Human *VDAC* pseudogenes: an emerging role for *VDAC1P8* pseudogene in acute myeloid leukemia

**DOI:** 10.1101/2023.01.20.524880

**Authors:** Xena Giada Pappalardo, Pierpaolo Risiglione, Federica Zinghirino, Angela Ostuni, Daniela Luciano, Faustino Bisaccia, Vito De Pinto, Francesca Guarino, Angela Messina

## Abstract

**Background:** Voltage-dependent anion selective channels (VDACs) are the most abundant mitochondrial outer membrane proteins, encoded in mammals by three genes, *VDAC1*, *2* and *3*, mostly ubiquitously expressed. As ‘mitochondrial gatekeepers’, *VDACs* control organelle and cell metabolism and are involved in many diseases. Despite the presence of numerous *VDAC* pseudogenes in the human genome, their significance and possible role in VDAC protein expression has not yet been considered.

**Results:** We investigated the relevance of processed pseudogenes of human *VDAC* genes, both in physiological and in pathological contexts. Using high-throughput tools and querying many genomic and transcriptomic databases, we show that some *VDAC* pseudogenes are transcribed in specific tissues and pathological contexts. The obtained experimental data confirm an association of the *VDAC1P8* pseudogene with acute myeloid leukemia (AML).

**Conclusions:** Our *in-silico* comparative analysis between the *VDAC1* gene and its *VDAC1P8* pseudogene, together with experimental data produced in AML cellular models, indicate a specific over-expression of the *VDAC1P8* pseudogene in AML, correlated with a downregulation of the parental *VDAC1* gene.

## Introduction

Voltage dependent anion selective channels (*VDACs or* mitochondrial porins) are a phylogenetically conserved family of genes encoding channel proteins embedded into the outer mitochondrial membrane (OMM). VDAC proteins are considered the master regulators of the mitochondrial function because the important functions they perform (1). The principal role of VDACs is to allow the exchange of important ions and metabolites to and from the mitochondria (2–4). In mammals, three *VDAC* isoforms have been identified and named *VDAC1, VDAC2* and *VDAC3* (5). VDACs genes are located in different autosomes but sharing the same number and size of exons. The only exception is *VDAC2* gene which contains an additional coding exon at the N-terminal (5,6). Also, the size of introns is variable (7). We recently reported that *VDAC* genes show a different expression pattern in human tissues, as well as a distinct set of transcription factor binding sites (TFBSs) on each promoter (8,9). Furthermore, it is known that altered *VDAC1* expression is associated with many diseases, like cancer, neurodegenerative and cardiovascular diseases (10–14). In particular, the observation that VDAC1 is over-expressed in most tumors suggests that this protein may play a pivotal role in this pathological condition (15). And concerning the tumor phenotype, the interaction of VDAC1 with hexokinase (HK), the first glycolysis enzyme, grants the enzyme immediate access to newly synthesized ATP and confers protection from apoptosis (16). It is therefore not surprising that VDAC1 has become a primary target in the fight against cancer.

In recent years, the involvement of transcribed pseudogenes in the development and progression of many types of cancer has become well established (17). Due to the vast amount of data available in the literature and in public databases, specific pseudogenes, whose expression varies considerably in certain cancers, can be classified as predictive, heritable or prognostic biomarkers.

Pseudogenes have long been considered function-less elements arisen during genome evolution as failed copies of genes. However, a large number of recent evidence indicate that most of them are not functionally inert but rather, once transcribed, have a consistent role in protein-coding genes regulation (18). Pseudogene expression pattern can vary under different pathological conditions including diabetes and cancer, even though, due to the high sequence similarity with the parental genes. However, measurements of their transcription level remain difficult to perform (19,20). Not surprisingly, considering the homology between pseudogene and its parental gene, pseudogene transcripts can play a part as competitive endogenous RNAs (ceRNAs). Generally, any transcript that competitively binds a common microRNA (miRNA) behaves as a ceRNA that, acting as a “sponge”, reduces the free miRNA available. As a result, this can lead to the release or attenuation of miRNA target gene(s) repression. Alternatively, RNA resulting from pseudogene transcription may be processed into a short interfering RNA (siRNA) or act as antisense RNA. Regardless of the mechanism, the pseudogene would in any case be an important factor in the regulation of protein-coding genes. Recently, it has been proposed that pseudogenes represent primate- or human-specific regulatory elements, especially in hemopoiesis, given their higher expression in bone marrow than in other tissues (21).

In the human genome there are 13 processed pseudogenes (or retrocopies) of *VDAC1*, distributed in 10 different chromosomes and annotated in GENECODE and NCBI Entrez Gene (22). Processed pseudogenes originate when a mRNA transcript is reverse-transcribed and integrated into the genome at a new position (23,24). Therefore, lacking its own promoter sequence and introns as well, their transcriptional activity depends on the presence of promoter(s) and regulatory sequences nearby. Ido and co-workers, after a first characterization of 16 putative rat *VDAC1* pseudogenes (25), reported a comparison of *VDAC1* pseudogenes among rat, mouse and human showing that no synteny exists between humans and rodents (26). Apart from these literature reports, nothing more is known about human *VDAC* pseudogenes. Considering the many regulatory roles described for pseudogenes in cancer (17,20,27) and the known dysregulation of VDAC in cancer pathologies (13,15,16), we wanted to investigate whether there is an association of any pathology, and in particular cancer, with *VDAC* pseudogenes expression. The assumption is that, in pathological situations, the expression of VDAC proteins could be post-transcriptionally regulated and/or influenced by the transcription of related pseudogenes. This information could be of great importance because it could certainly be useful in identifying new diagnostic, prognostic and therapeutic approaches for the pathologies under consideration.

In this work, we present a whole, comprehensive analysis of in-silico data concerning the human *VDAC* pseudogenes together with a careful *in-cellulo* validation of the expression data. Overall, our results allow us to hypothesize for some *VDAC1* pseudogenes a role in the physiology or pathology of certain tissues. In particular, the specific and marked expression in AML of the *VDAC1P8* pseudogene suggests its potential role in leukaemogenesis and makes it a possible biomarker of acute myeloid leukaemia.

## Results

### Human VDACs pseudogenes and their relative chromatin state

All the genomic databases (NCBI Entrez Gene, UCSC and Ensembl) queried in this work report the presence in the human genome of 13 *VDAC1* pseudogenes, 4 *VDAC2* pseudogenes and only one *VDAC3* pseudogene. Evidence in human genome of putative regulatory elements located in the proximity of *VDAC* pseudogenes was obtained from UCSC Genome Browser (GRCh38/hg38).

The data on regulatory elements found for each *VDAC* pseudogene are summarized in Table 1. This analysis revealed that *VDAC1P4*, *VDAC1P8*, *VDAC1P11* and *VDAC1P13* pseudogenes are located at transcriptionally active genomic loci (Table 1). Interestingly, the *VDAC1P8* sequence is located between the 5’ end of the *PEX3* gene and the 3’ end of the *FUCA2* gene, on the opposite strand (Table 1, Fig.1). Therefore, *VDAC1P8* gene sequence is found within a likely active region characterized by lax chromatin states, the presence of strong and weak enhancers and a weak transcriptional elongation capacity (ENCODE genome segmentation). Although the state of the chromatin is not comparable to that in which housekeeping genes, as GAPDH, are located, the features of chromatin region hosting *VDAC1P8* pseudogene suggest the possibility of transcriptional activation of this genomic stretch, unlike the chromatin region where *VDAC1P5* is found (Fig.1, Suppl. Fig.2, Suppl. Fig.3). Although the genomic regions of the pseudogenes *VDAC1P3*, *VDAC1P4*, *VDAC1P11* and *VDAC1P13* are associated with moderate transcriptional activity, their sequences overlap with introns of other genes (*PCDH11-X, ACBD6, ZNF169* and *KCNG-3*, respectively) apparently unrelated to *VDAC*. Their localization suggests that, in the absence of a specific maintenance mechanism, the fate of these sequences is to be degraded once spliced. The *VDAC1P1*, *VDAC1P6*, *VDAC1P7*, *VDAC1P9* and *VDAC1P10* sequences are instead associated with heterochromatin, reduced transcriptional activity and weak enhancers (Table 1). Likewise, the remaining *VDAC1* pseudogenes (*VDAC1P2*, *VDAC1P5*, *VDAC1P12*) along with all four *VDAC2* pseudogenes and the only one from *VDAC3* are located near transcriptionally inactive domains and associated with polycomb repressors (Table 1).

**Table 1.** Features and chromatin status of genomic sequences containing *VDAC* pseudogenes, from UCSC Genome Browser (GRCh38/hg38). Table shows in (a) information on *VDAC1* pseudogenes (*VDAC1P1-13*), in (b) on *VDAC2* pseudogenes (*VDAC2P1-4*) and in (c) about the only one *VDAC3* pseudogene (*VDAC3P1*). *\*Genomic position is referred to splicing variant with ENST00000406025.2 ID at Ensembl database (Suppl. Fig.1a).*

| (a) |  | *Genomic Coordinates* |  |  | *Regulatory Build* |  |
| --- | --- | --- | --- | --- | --- | --- |
| Name | Entrez ID | Location | Size (bp) | Strand | 5'-flanking region | 3'-flanking region |
| ***VDAC1P1*** | 642585 | chrX:80184923-80186706 | 1784 | + | heterochromatin | insulator and weak enhancer |
| ***VDAC1P2*** | 10064 | chrX:49396238-49397962 | 1725 | - | heterochromatin |  |
| ***VDAC1P3*** | 100310838 | chrX:91236801-91238513 | 1713 | - | weak enhancers, weak and active promoter (overlapping intronic region of *PCDH11X* gene) |  |
| ***VDAC1P4*** | 7418 | chr1:180403931-180405937 | 2007 | + | weak and strong enhancers, weak transcriptional elongation, weak promoter (overlapping intronic region of *ACBD6* gene) |  |
| ***VDAC1P5*** | 10187 | chr12:55195669-55197407 | 1739 | - | heterochromatin |  |
| **VDAC1P6** | 359800 | chrY:5074406-5076119 | 1714 | - | heterochromatin (overlapping intronic region of *PCDH11Y* gene) |  |
| **VDAC1P7** | 100310839 | chr3:77365047-77366767 | 1721 | - | insulator | heterochromatin (overlapping intronic region of *ROBO2* gene) |
| **VDAC1P8\*** | 100310840 | chr6:143811645-143813335 | 1691 | + | weak enhancer, weak transcriptional elongation (proximity to *PEX3* gene) | weak and strong enhancer, weak transcriptional elongation (proximity to *FUCA2* gene) |
| **VDAC1P9** | 391106 | chr1:157693990-157694816 | 827 | - | heterochromatin | weak enhancer, weak transcriptional elongation |
| **VDAC1P10** | 643536 | chr1:215549841-215551581 | 1741 | + | flanking heterochromatin and weak enhancers inside gene body |  |
| **VDAC1P11** | 100310841 | chr9:97048814-97050837 | 2024 | - | weak transcriptional elongation, weak and strong enhancer (overlapping intronic region of *ZNF169* gene) |  |
| **VDAC1P12** | 100874289 | chr13:34656914-34657447 | 534 | - | heterochromatin |  |
| **VDAC1P13** | 100420568 | chr2:42690279-42691137 | 859 | - | weak enhancers, weak transcriptional elongation (overlapping intronic region of *KCNG3* gene) |  |

| (b) |  | **Genomic Coordinates** |  |  | **Regulatory Build** |  |
| --- | --- | --- | --- | --- | --- | --- |
| Name | Entrez | Location | Size (bp) | Strand | 5'-flanking region | 3'-flanking region |
| **VDAC2P1** | 54015 | chr21:17466509-17467792 | 1284 | - | heterochromatin (overlapping *LINC00478* gene) |  |
| **VDAC2P2** | 643996 | chr12:9188920-9190119 | 1200 | - | heterochromatin |  |
| **VDAC2P3** | 401959 | chr1:118182962-118184349 | 1388 | - | insulator, weak enhancer, heterochromatin | weak transcriptional elongation, polycomb repressed, heterochromatin |
| **VDAC2P4** | 100420574 | chr2:135554739-135555449 | 711 | + | heterochromatin, insulator, Polycomb repressor |  |

**Table 1. Features and chromatin status of genomic sequences containing *VDAC* pseudogenes, from UCSC Genome Browser (GRCh38/hg38).**
| (c) |  | **Genomic Coordinates** |  |  | **Regulatory Build** |  |
| --- | --- | --- | --- | --- | --- | --- |
| Name | Entrez ID | Location | Size (bp) | Strand | 5'-flanking region | 3'-flanking region |
| ***VDAC3P1*** | 341965 | chr14:100430801-100431912 | 1112 | + | weak transcriptional elongation, heterochromatin, Polycomb repressed | weak enhancer, heterochromatin, Polycomb repressed |

**Figure 1.**
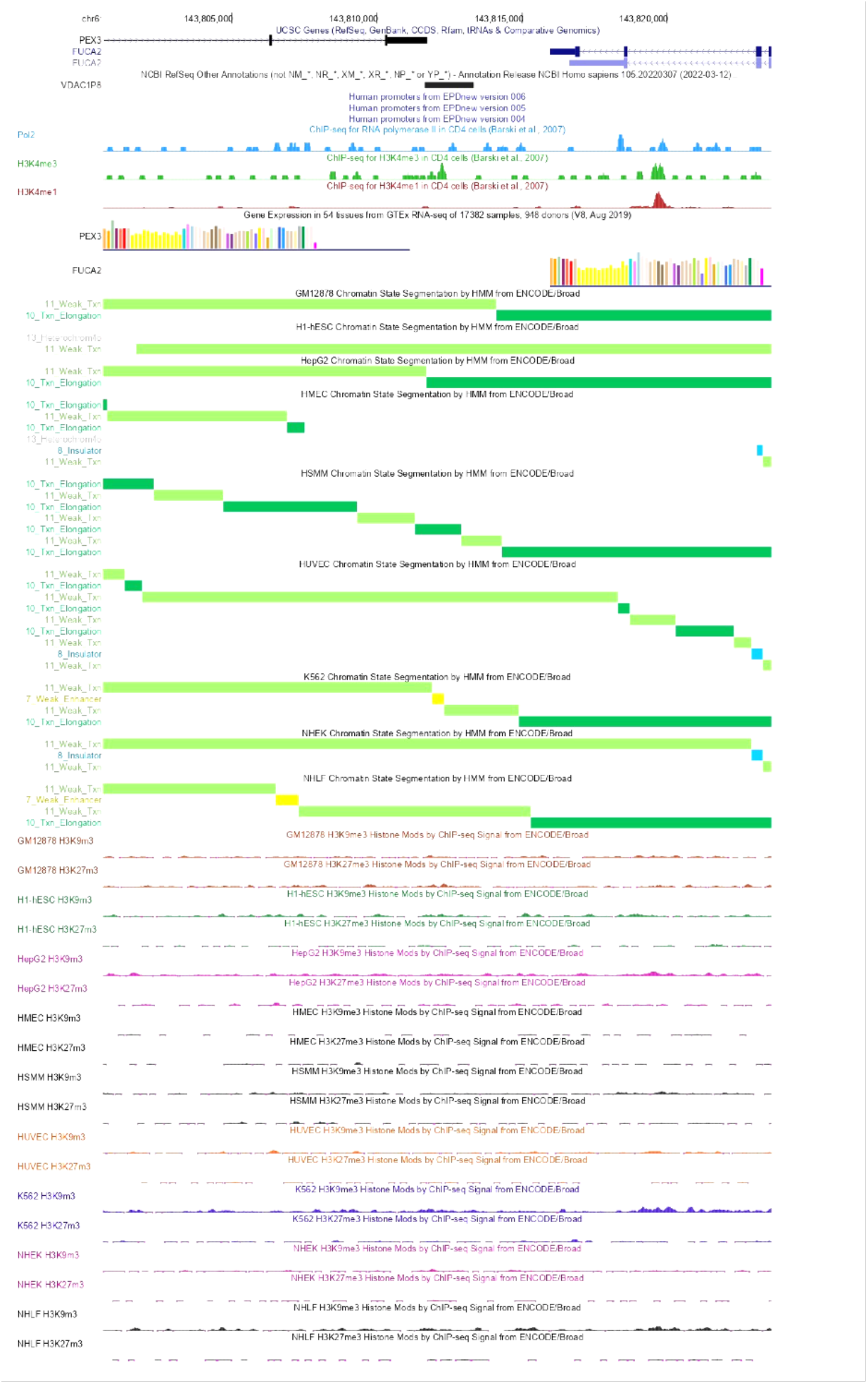
Chromatin state and genomic features of *VDAC1P8* pseudogene from UCSC Genome Browser GRCh37/hg19. The genomic context of *VDAC1P8* pseudogene set around 1000 Kb upstream and downstream of the annotated Refseq is shown. The selected regulatory hub tracks are Pseudogene Annotation Set from GENCODE v.38lift37 Ensemble 104, Eukaryotic Promoter Database EPD v.4-6, CpG island track, Genotype-Tissue Expression GTEx RNA-seq v.8 2019, ChIP-Seq data for RNA polymerase II, H3K4me3 and H3K4me1, used as markers of transcriptional activation, while H3K9me3 and H3K27me3 are markers of transcriptional repression, and chromatin state segmentation by Hidden Markov Model from the ENCODE/Broad project of nine different cell lines (GM12878, H1-hESC, HepG2, HMEC, HUVEC, K562, NHEK, NHLF) identified using the following different colors: yellow=weak/poor enhancer; blue= isolator; dark green= transcriptional transition/elongation; light green= transcriptional transcript.

### Expression of *VDAC* genes and their pseudogenes in tumour and normal tissues

To find support for the chromatin status analysis data and those obtained by querying UCSC Genome Browser, we analysed the transcriptomic data of *VDAC* genes (Suppl. Table 1) and their pseudogenes (Suppl. Table 2) released from the GEPIA repository. A comparative analysis of expression profiles in tumour and normal samples was performed for most of the *VDAC1* pseudogenes. As suggested by the active chromatin status where *VDAC1P8* is located, we found that this pseudogene is widely expressed in both normal and tumour tissues, with an expression value in the range of 1 to 9 transcript per million (TPM) in normal tissues, with the exception of a few cases (Suppl. Table 2). In particular, the highest expression level is reported in uterus, ovary, lung, rectum, colon, thyroid. Also, *VDAC1P8* is downregulated in almost all tumour tissues considered, compared with its normal counterpart. An exception is acute myeloid leukemia (AML) where, surprisingly, *VDAC1P8* is overexpressed with a TPM value reaching the highest value of 17.75 TPM (2.5 times higher than normal tissue). *VDAC1P1* is also a pseudogene expressed in all tissues and their tumour counterpart, in most case at comparable levels and in the range of 0.01-0.9 TPM (Suppl. Table 2). The *VDAC1P2* pseudogene is expressed, with the exception of AML, in all tumours examined, at levels between 0.11 and 0.44 TPM in the stomach and kidney, respectively. In addition, *VDAC1P2* is expressed in some normal tissues, at levels between 0.01 and 0.26 TPM. (Suppl. Table 2). Similarly, the *VDAC1P6* pseudogene, located in a region of repressed chromatin is expressed in the range of 0.01-0.07 TPM in normal tissues (head, neck, kidney, bone marrow, bone and “soft” tissues - fat, muscle, nerves) and in their tumour counterparts with the highest value in colon and rectum adenocarcinoma (Suppl. Table 2).

Also, by examining the data retrieved from the GEPIA server, we determined that the pseudogenes *VDAC1P3, VDAC1P4, VDAC1P9, VDAC1P13, and VDAC1P11* are expressed in only very few tissues and at very low levels (0.01-0.1 TPM). No expression data were found for the pseudogenes *VDAC1P5, VDAC1P7, VDAC1P10, VDAC1P12* and *VDAC2P2-4*. Indeed, these pseudogenes are located in regions of repressed chromatin or where only weak active chromatin elements are present.

Furthermore, we observed that *VDAC2P1* and *VDAC3P1* are weakly expressed (range of 0.03-0.07 TPM) only in some normal connective tissues (i.e. tissues subject to sarcomas), although *VDAC3P1* is also expressed in testicular germ cells (Suppl. Table 2).

Our search in the GEPIA server for the expression profiles of *VDAC1-3* genes provided us with useful information as to whether the transcription of active *VDAC* pseudogenes could be linked to the expression of their parental genes. Exactly, this analysis showed that all three *VDAC* isoforms are overexpressed in many tumors, with the exception of AML, where their transcriptional levels were lower than in healthy tissue (Suppl. Table 1). Focusing only on expression alterations in AML, we have collected in Fig. 2a-b the results of the comparative analysis of the transcriptional profiles of *VDAC1* gene and its pseudogenes specifically reported for AML and the normal counterpart. In particular, we observed a correlation in AML between the significant downregulation of the *VDAC1* gene (from 221.12 TPM in normal tissue to 80.01 TPM in tumour) in AML and the expression levels of some pseudogenes, especially *VDAC1P8* pseudogene. Indeed, *VDAC1P1* and *VDAC1P2* show a reduction in AML, while *VDAC1P4* and *VDAC1P11* an increase over their normal counterpart but with expression levels below 1 TPM. An exception is *VDAC1P11*, which shows a change of 1.69 TPM units in AML compared with normal tissue. Interestingly, the expression level of *VDAC1P8* in AML increases of 2.5 folds, reaching the value of 17.50 TPM compared to its normal counterpart of 7.10 TPM. These intriguing data prompted us to pay attention to the putative role of *VDAC1* pseudogenes in AML and, in particular, to further investigate the involvement of the *VDAC1* gene/pseudogene *VDAC1P8* pair.

**Figure 2a-d.**
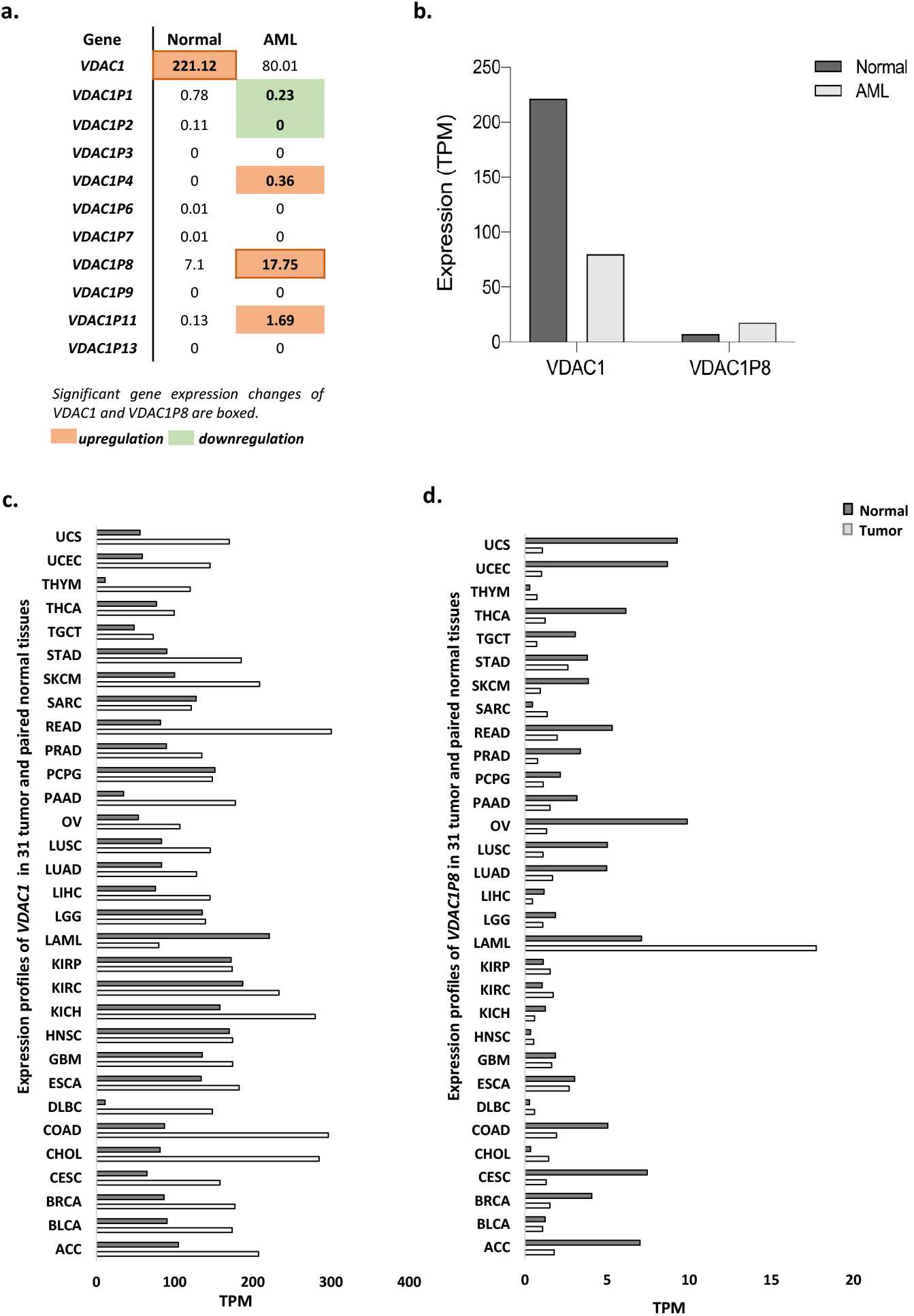
Analysis of gene expression profiles of *VDAC1* and its pseudogenes from GEPIA repository. Specific-AML expression changes of *VDAC1* compared to its pseudogenes was shown (a). Only values for *VDAC1* and *VDAC1P8* are plotted (b). Gene expression trend changes in tumor lines of *VDAC1* and *VDAC1P8* and in their normal counterparts are graphically represented (c-d). Data shown are included in Supp. Tables 1-2. The height of bars is the median expression of Log2TPM+1 of normal and AML tissue based on GTEX and TCGA respectively, available from GEPIA repository. TPM= transcripts per kilobase million.

Indeed, the expression pattern of *VDAC1* gene in tumours revealed that AML is the only type in which this gene is exclusively downregulated. On the contrary, it is even more interesting to note that in AML there is a significant increase in *VDAC1P8* pseudogene expression (Fig.2c-d).

### The putative involvement of 6q24.2 locus in AML

AML is a common haematological disorder with abnormal proliferation and differentiation of immature myeloid cells, characterised by genetic and clinical heterogeneity and high mortality (28). Despite the development of new therapies, most patients with AML continue to relapse, highlighting the existence of an unresolved problem. More recently, the role of non-coding RNAs (ncRNAs) in the biology of AML and in the mechanisms of resistance to therapy has begun to be explored (29). The full potential of circular RNA (circRNA), miRNA and long noncoding RNA (lncRNA) as diagnostic, therapeutic and prognostic factors in AML is thus emerging (30–32). Also, the involvement of specific pseudogenes is beginning to appear. Our data therefore point in this direction. In particular, they show that, interestingly, AML is the only tumor type in which the *VDAC1* gene is downregulated and its *VDAC1P8* pseudogene is clearly up-regulated (Fig.2a-b).

Furthermore, since *VDAC1P8* pseudogene is over-expressed exclusively in AML (Fig.2a-b) we then wished to investigate whether other genes located in the same genomic locus and putatively under the effect of the same regulatory elements are also associated with the same disease. By querying the GeneCards database (https://www.genecards.org/) we found that the best enhancer associated with *VDAC1P8* also controls the expression of 13 other target genes (*ADAT2*, *PEX3*, *HSALNG0054032*, *RF00001-290*, *RNA5SP221*, *LTV1*, *ENSG00000270890*, *PLAGL1*, *TUBB8P2*, *ENSG00000278206*, CM03496-315, *FUCA2*, *MN298114-190*), located in the same 6q24.2 cytogenetic region. Interestingly, 5 genes (*ADAT2*, *ENSG000278206*, CM03496-315, *FUCA2*, *MN298114-190*) as *VDAC1P8* are also correlated with AML (Suppl. Table 3).

These results were confirmed by querying the GEPIA database from which we also extrapolated altered expression data in AML for the *ENSG00000270890*, *TUBB8P2* and *LTV1* genes. Thus, the 6q24.2 locus would appear to have a strong association with AML. It is also noteworthy that as many as 7/13 genes (*RNA5SP221*, *HSALNG0054032*, *ENSG00000270890*, *TUBB8P2*, *ENSG00000278206*, CM03496-315, *MN298114-216*) with the same cytogenetic localization of *VDAC1P8* pseudogene produce ncRNA (3 pseudogenes and 4 lncRNAs). Unfortunately, there is little or no existing expression data for these genes.

### Sequence analysis of *VDAC1* pseudogenes transcripts

From an initial analysis on human *VDAC* genes and their respective pseudogenes, we noticed an interesting downregulation in AML of the *VDAC1* gene, compared to its expression in normal tissue, associated with increased expression of the *VDAC1P8, VDAC1P1* and *VDAC1P11* pseudogenes. This prompted us to study in more detail the relationships between the *VDAC1* gene and its pseudogenes. We thus proceeded to analyze the sequences of the *VDAC1* pseudogenes by comparing them with the transcript of the parental gene. The multialignment shows the high homology between the pseudogenes and *VDAC1* sequences (Suppl. Fig. 4a), in particular the *VDAC1P1* transcript matches exactly the *VDAC1* ORF with which it has a very high sequence homology (97%). Multi-alignment also reveals the existence of additional sequence segments to the *VDAC1* ORF in only a few pseudogenes (*VDAC1P10, VDAC1P12, VDAC1P13*). The *VDAC1P8* pseudogene was not included in this multi-alignment analysis because, unlike other *VDAC1* pseudogenes that were assigned a single processed transcript, *VDAC1P8* has multiple transcripts (Suppl. Fig.4b). Indeed, fourteen non-coding splicing variants, classified according to their biotypes, are annotated for this *VDAC1* pseudogene in GENCODE v.39 (Ensembl 105) (Suppl. Fig. 5a), as in all other databases interrogated. Among these transcript variants, 8/14 variants are reported as “intron retained”, 5/14 as “processed transcripts” and only one as a “transcribed processed pseudogene”. By analyzing the sequences of the *VDAC1P8* splice variants, emerges that the *VDAC1P8-201* transcript consists of a single exon aligning almost perfectly (89% homology) with *VDAC1* ORF sequence (Suppl. Fig.4b, Suppl. Fig. 5). In contrast, *VDAC1P8-210* and *VDAC1P8-205* transcripts contain 5 and 4 exons, respectively, of which only the first exon overlaps with the *VDAC1P8-201* exon (87.53% and 89.28% sequence homology between the *VDAC1* ORF and the 1st exon of *VDAC1P8-205* and *VDAC1P8-210*, respectively), whereas the other exons have unrelated sequences to the parental *VDAC1* gene (Suppl. Fig. 4b, Suppl. Fig. 5). Similarly, the remaining alternative transcripts from *VDAC1P8* contain exon sequences that are not shared with the *VDAC1* ORF, so they should not be considered as correlated to this gene (Suppl. Fig.4b, Suppl. Fig.5). Later in the text, the term *VDAC1P8* is used to refer to the −201 variant of this pseudogene, unless otherwise specified.

### Search for orthologs of *VDAC1* pseudogenes in the highest primates

Literature (5,6,25,25) combined with data in the NCBI Entrez Gene database (https://www.ncbi.nlm.nih.gov/gene/?Term=related_functional_gene_22333%5Bgroup%5D) show that several *VDAC* pseudogenes are predicted in the *Mus musculus* genome, exactly: thirteen for *mVDAC1*, three for *mVDAC2* and three more for *mVDAC3*. In their work, Ido et al. (25,26) suggest the existence of syntenic relationships in the mouse and rat genomes between two putative rat *VDAC1* pseudogenes and one mouse *VDAC1* pseudogene, respectively. These authors also point out the lack of synteny between rodent *VDAC1* pseudogenes and humans. However, apart from the results from this outdated study focused mainly on rodent *VDAC* pseudogenes, nothing to date is known about the presence of homologs of *VDAC1* pseudogenes in higher primates.

To compensate for the lack of this and other important information, by using the Ensembl BLASTN tool we searched primate genomes for the presence of orthologous genes to the 13 human *VDAC1* pseudogenes. Sequences homologous to *VDAC1* pseudogenes were identified in all genomes examined (9 primate species, belonging to different evolutionary groups: Marmoset, Macaque, Drill, Baboon, Gibbon, Orangutan, Gorilla, Bonobo, Chimpanzee). Unfortunately, the sequences identified mostly fall into genomic tracts that are either unannotated or overlapping with coding gene sequences or, in some cases, correspond to genes for ncRNAs that have not yet been characterized. Therefore, no firm data were found on the possible expression of orthologs of the *VDAC1* pseudogenes. This lack of information combined with the high homology of the 13 query sequences used in the BLAST analysis led to the identification in several primate species of the same sequence as homologous to various human pseudogenes of *VDAC1*. For these reasons but also considering the specific overexpression of *VDAC1P8* in AML, we restricted the homology analysis to this pseudogene only. This BLAST analysis showed that in all primate species examined there is a sequence highly homologous to the human *VDAC1P8* pseudogene, which is very well maintained starting from new world monkeys to hominids (Suppl. Table 4).

Then, starting from the genomic location of each *VDAC1P8* ortholog and using Ensemble’s “Synteny” tool, we assessed whether each genomic locus fell within the known synteny blocks between human and other primate species’ chromosomes. We thus found that in all primate species analyzed, the orthologs of *VDAC1P8* are maintained within the same genomic synteny blocks with humans (Suppl. Table 4). These genomic regions correspond to stretches of active chromatin in which the other genes associated with the human 6q24.2 locus and shown in section 3.3 are also retained. The presence in all higher primates of sequences highly homologous to *VDAC1P8* in potentially expressed genomic regions allows us to hypothesize an important role for this backcopying, already in the physiological context.

### Expression patterns and methylation profile of *VDAC1* gene and *VDAC1P8* pseudogene in healthy adult and fetal human tissues

The expression profiles data of the *VDAC1* gene and its pseudogenes, in particular *VDAC1P8*, obtained by querying the GEPIA server prompted us to further investigate their physiological expression in human adult and fetal tissues. For this aim, we analyzed the RNA-Seq CAGE (Cap Analysis of Gene Expression) RIKEN FANTOM5 project from Expression Atlas (www.ebi.ac.uk/gxa/home). Thus, we confirmed that *VDAC1* gene and *VDAC1P8* pseudogene are both normally expressed in any healthy adult and fetal tissues reported. The expression level of the *VDAC1* gene is comparable between adult and fetal tissues while the *VDAC1P8* pseudogene is more expressed in fetus (Fig. 3b). Exactly, in adult tissues the highest expression levels of *VDAC1P8* are found in bone marrow, diencephalon and heart (Fig. 3a); whereas in the fetus, there is an abundant expression in the colon, duodenum, small intestine and throat, which is up to ten times higher than *VDAC1* gene transcript (Fig. 3b). Therefore, *VDAC1P8* is mainly expressed in fetal tissues, although it is during embryogenesis that pseudogenes silencing generally occurs (33). In particular, by performing a correlation analysis of the expression levels of *VDAC1* and its pseudogene *VDAC1P8* in the reported tissues, it appears that adult and fetal tissues with subthreshold expression of *VDAC1P8* have high levels of *VDAC1*. Likewise, tissues with high levels of *VDAC1P8* expression have reduced levels of *VDAC1* (with the exception of bone marrow in adults). No data were available in the database for the other pseudogenes of *VDAC1*.

**Figure 3a-b.**
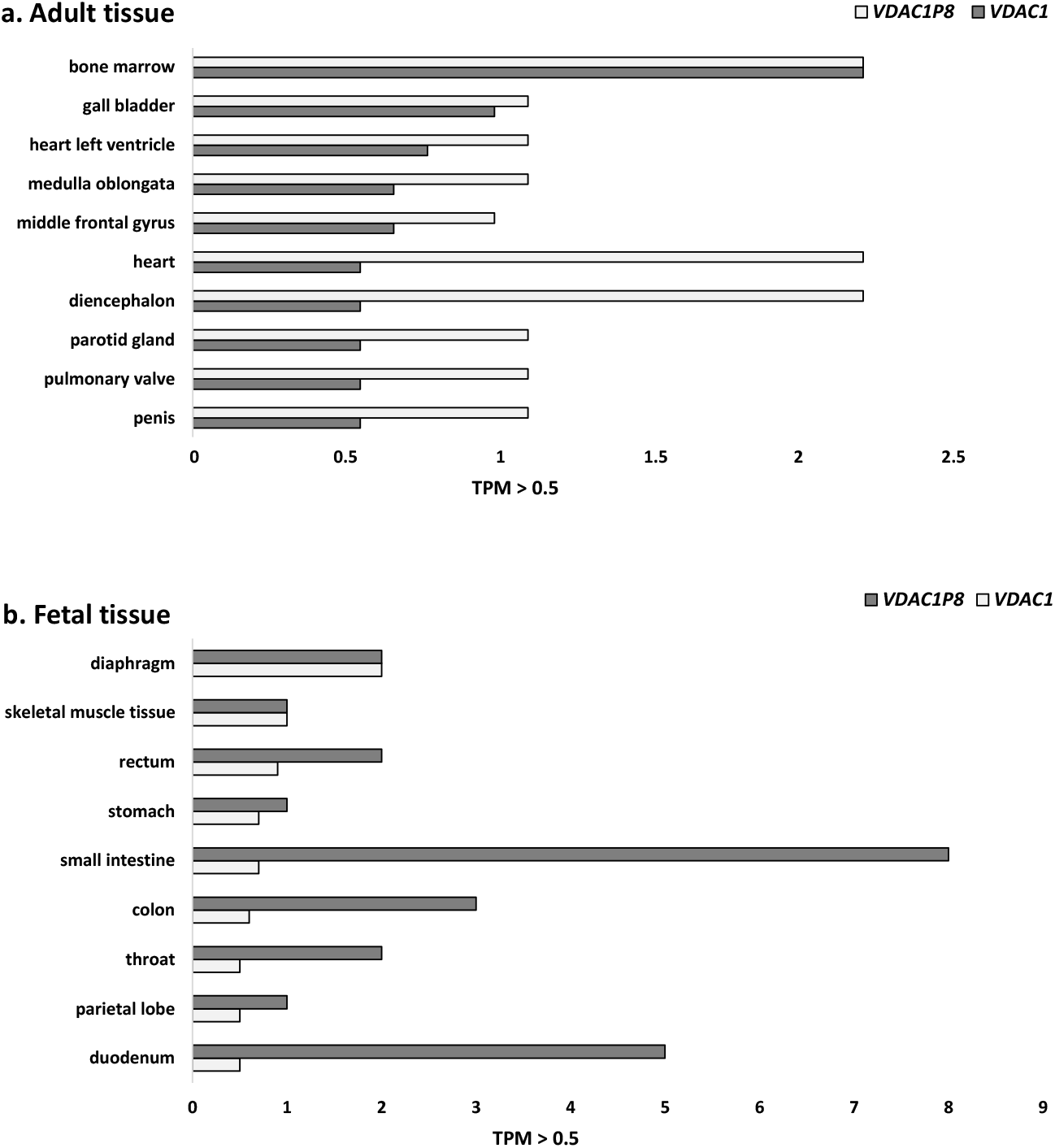
Adult (a) and fetal (b) transcripts tissue expression levels of *VDAC1* and *VDAC1P8* from RNA-seq CAGE in RIKEN FANTOM 5 project. Data cut-off is >0.05 TPM. Data shown consist of RNA-Seq CAGE (Cap Analysis of Gene Expression) analysis from RIKEN FANTOM5 project (www.ebi.ac.uk/gxa/home). TPM= transcripts per kilobase million.

Furthermore, the methylation profile of putative promoter sequences and gene body of *VDAC1* gene and *VDAC1P8* pseudogene was investigated across the results retrieved from MethBank. As resumed in the Fig.4a-b, the β-value expressing the methylation level is clearly higher in the promoter region and in the gene body of *VDAC1P8* pseudogene in comparison to *VDAC1* gene for all the samples tissue derived from the central nervous system, circulation system, digestive system and lymphatic system. In particular, the promoter region of *VDAC1P8* presents a degree of methylation which is about 10 folds higher than *VDAC1* gene. The methylation profile of the other *VDAC1* pseudogenes regulated in AML is similar to that of *VDAC1P8* (Suppl. Fig.6a-b). Altogether, the information available on methylation profile of *VDAC* genes are very poor. However, the level of CG methylation of the human *VDAC* genes revealed that, in human and in mouse, the *VDAC1* promoter is less methylated than the other two isoforms confirming its ubiquitous and stable expression as housekeeping gene. The methylation profile of *VDAC1P8* pseudogene promoter suggests a control mechanism for its transcriptional activity which could be triggered in specific pathological conditions as in AML.

**Figure 4a-b.**
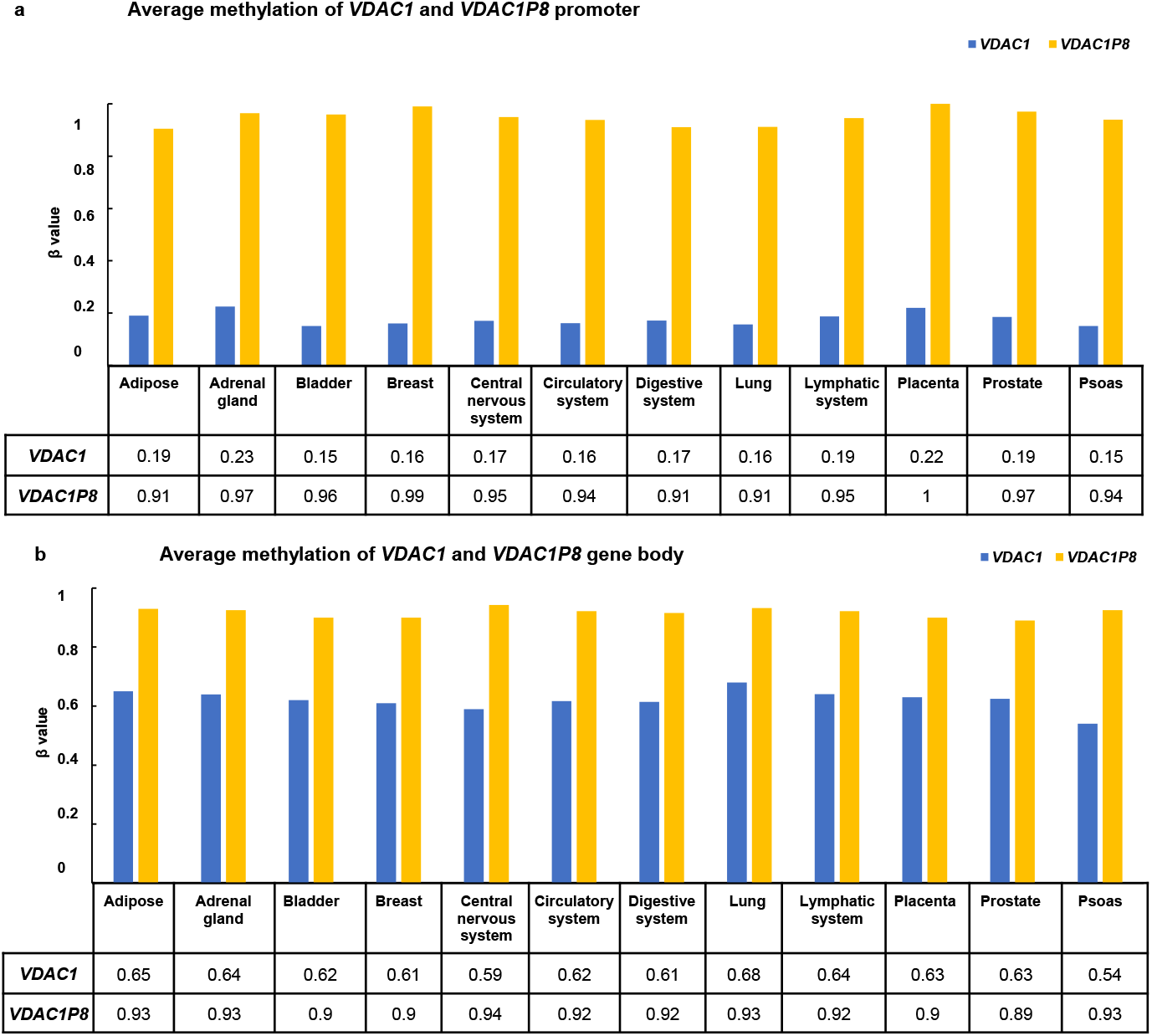
Methylation levels of *VDAC1P* and *VDAC1P8* gene in putative promoter (a) and gene body (b).* *The average methylation of* single-based resolution methylomes (*SRMs) data* provided by MethBank v.4.1 (https://ngdc.cncb.ac.cn/methbank) *are calculated as β-Value that reflects the methylation intensity at each CpG site. β-Values of 0–1 (represented from 0 to 1) indicate signifying percent methylation, from 0 to 100%, respectively, for each CpG site*.

### Putative TFBS suitable for driving *VDAC1P8* expression in leukemogenesis

The results of the expression analyses of *VDAC* pseudogenes showed that the pseudogene*s VDAC1P8*, *VDAC1P11 and VDAC1P4* are over-expressed in AML, whereas *VDAC1P1* and *VDAC1P2* are under-expressed, relative to their level in the corresponding normal tissues. We therefore searched for further evidence to support these gene expression data. Specifically, we analyzed the putative TFBS pattern of the *VDAC1* pseudogenes and compared it with that of the *VDAC1* gene. This analysis was performed by querying the hTFtarget database (http://bioinfo.life.hust.edu.cn/hTFtarget#!/) and selecting each candidate TF in blood and bone marrow. Interestingly, only the pseudogenes *VDAC1P8* and *VDAC1P11*, upregulated in AML, have binding sites for transcription factors involved in development of the tumor counterpart. We identified 24 TFBSs in the target regulatory region of *VDAC1P8* (Table 2), some of which (*MYC*, *TAL1*, *JUND*, *BATF*, *FOS*, *CEBPB/D*, *GATA1/2/3*, *CTCF*, *ZNF384*, *RCOR1*, *SPI1*, *ETS1*, *ERG*, *FLI1*, *IRF4* and *RUNX1*) had already been associated with leukemogenesis (34–49), while for others (*BRD4, CBX3, POLR2A*, *RAD21* and *STAG1*) the role of epigenetic modulators of the hematopoietic system, from normal to disease state, has only recently been demonstrated (38,50–52). In the regulatory region of *VDAC1P11*, six TFBS associated with leukemogenesis (*JUND, CBX3, RCOR1, GATA1, JUN, FOS*), have identified as possible regulators. To understand whether a common or differentiated regulatory pattern for the *VDAC1* gene and its pseudogenes *VDAC1P8* and *VDAC1P11* is associated with blood and bone marrow tumor development, the distribution of selected TFBS was compared (Fig. 5). The results clearly show that 19 of 24 TFBS found in the *VDAC1P8* regulatory region and involved in leukemogenesis are also putative regulators of *VDAC1* gene expression; while, only 3 of 8 were counted for *VDAC1P11*. Among all the transcription factors found, *JUND, CBX3, RCOR1* and *GATA1* are regulators commonly shared by the *VDAC1* gene and both *VDAC1P8* and *VDAC1P11* pseudogenes. In contrast, *VDAC1P1* appears to be associated with lamin B1 (*LMNB1*) transcription factor, which is known to be overexpressed in several cancers, such as lung adenocarcinoma, breast cancer and leukemia (53). Overall, these results suggest that *VDAC1* and the pseudogenes *VDAC1P8* and *VDAC1P11* share a common transcriptional regulatory pattern in the onset of leukemia.

**Table 2:**
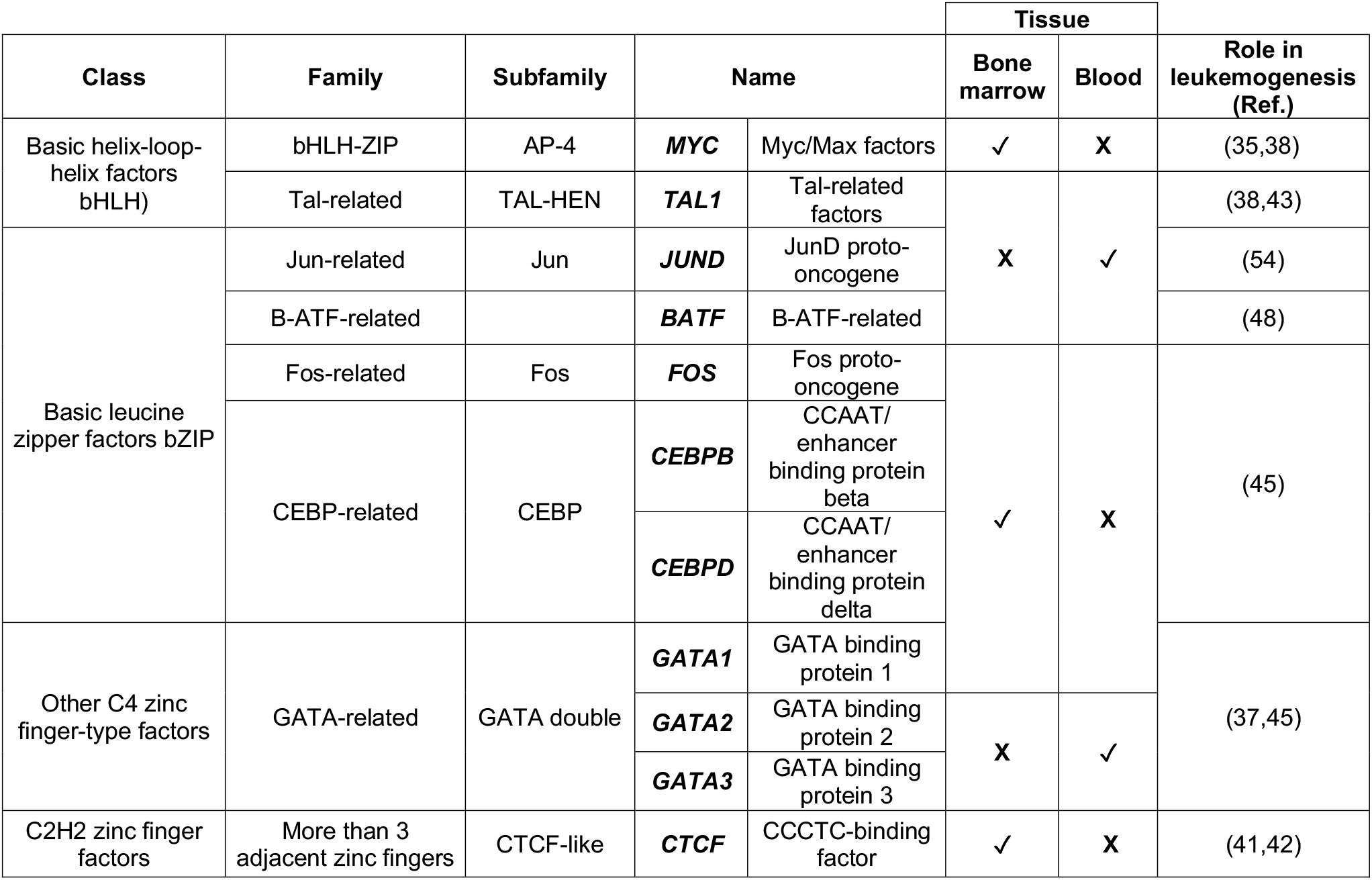
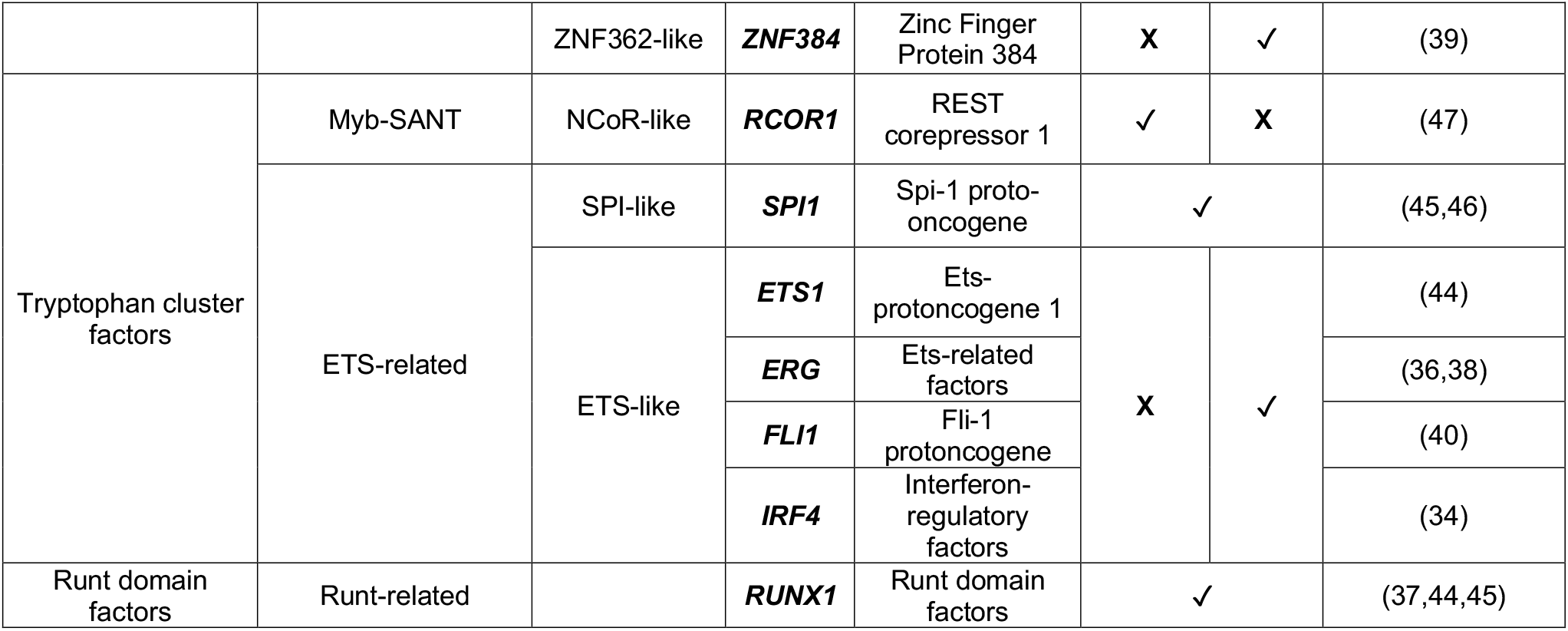
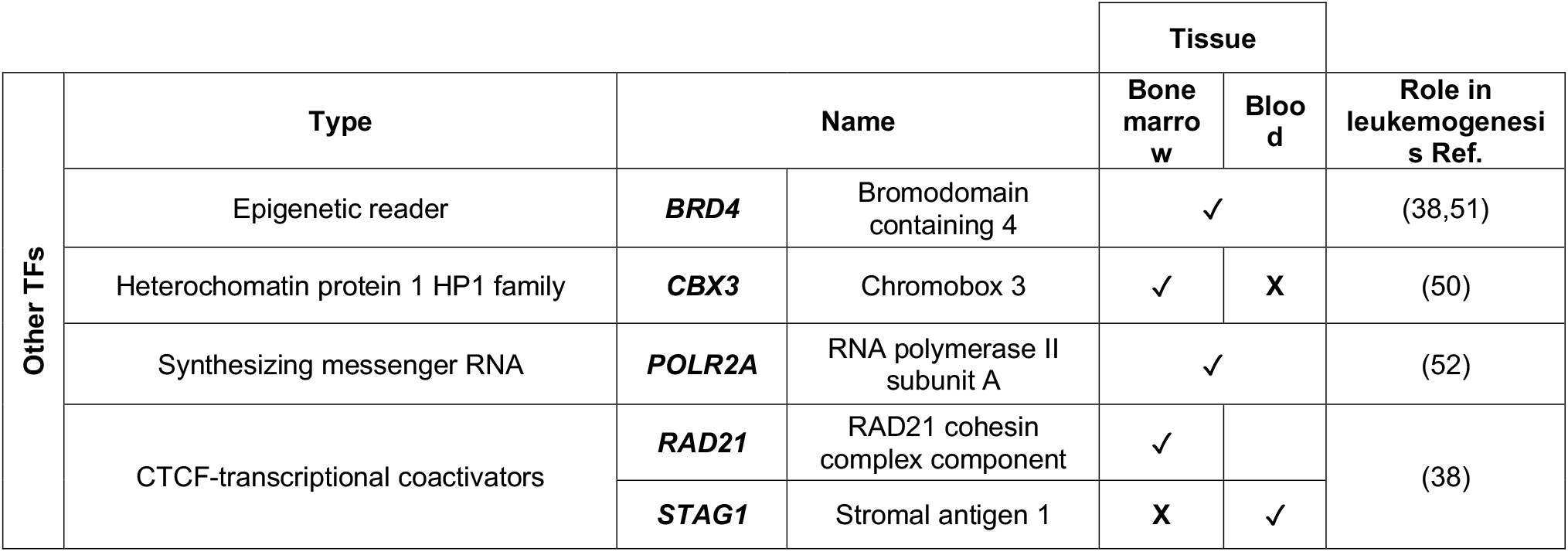
List of transcription factors (TFs) for *VDAC1P8* detected in blood and bone marrow. The analysis was performed by querying the hTFtarget database (http://bioinfo.life.hust.edu.cn/hTFtarget#!/) and selecting only candidate TFs found in blood and bone marrow.

**Figure 5a-d.**
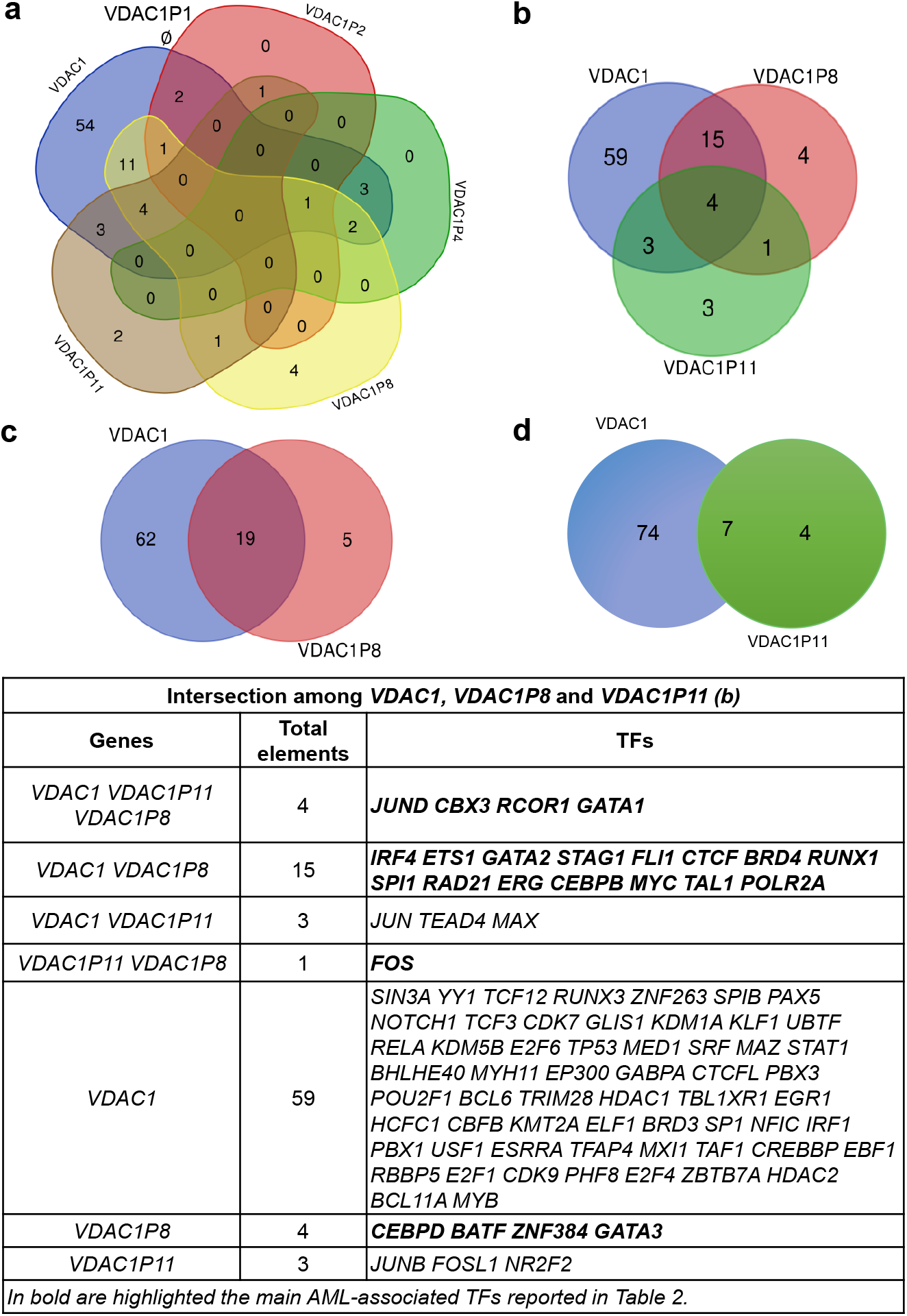
Comparative transcriptional analysis between *VDAC1* and its pseudogenes. The analysis of TFs was performed by querying the hTFtarget database (http://bioinfo.life.hust.edu.cn/hTFtarget#!/) selecting only candidate TFs found in blood and bone marrow. In (a), the Venn diagram shows the intersection among *VDAC1, VDAC1P2-P4-P8 and P11* with a null intersection of *VDAC1P1*. According to hTFtarget database, the TF found in *VDAC1P1* is lamin B1 (*LMNB1*), not shared by other pseudogenes. In (b), it is illustrated the intersection among *VDAC1, VDAC1P8* and *VDAC1P11*, which elements are represented in the table. In (c), the intersection between *VDAC1* and *VDAC1P8*. In (d), the intersection between *VDAC1* and *VDAC1P11*. The Venn diagrams made by https://bioinformatics.psb.ugent.be/cgi-bin/liste/Venn/calculate_venn.htpl.

### Expression of *VDAC1P8* pseudogene in leukemia cell lines

In light of the expression data of the *VDAC1* gene and its pseudogene *VDAC1P8* retrieved from public repository, we decided to experimentally evaluate their expression in 5 different acute myeloid leukemia cell lines. Exactly, we tested HL60, OCI-AML/2 and OCI-AML/3 (as myelocytic AML cell lines), MOLM13 (as monocytic AML cell line) and IMS-M2 (as megakaryocytic AML cell line) (55–57)

Confirming the data obtained in silico, both *VDAC1* and *VDAC1P8* were found to be expressed in all cell lines tested, with a higher transcript level of *VDAC1* than pseudogene (Fig.6a). By sequencing the amplification products obtained by the specific primers pairs (Suppl. Fig. 5), we confirmed the identity of *VDAC1-P8* pseudogene and *VDAC1* gene.

**Figure 6a-b.**
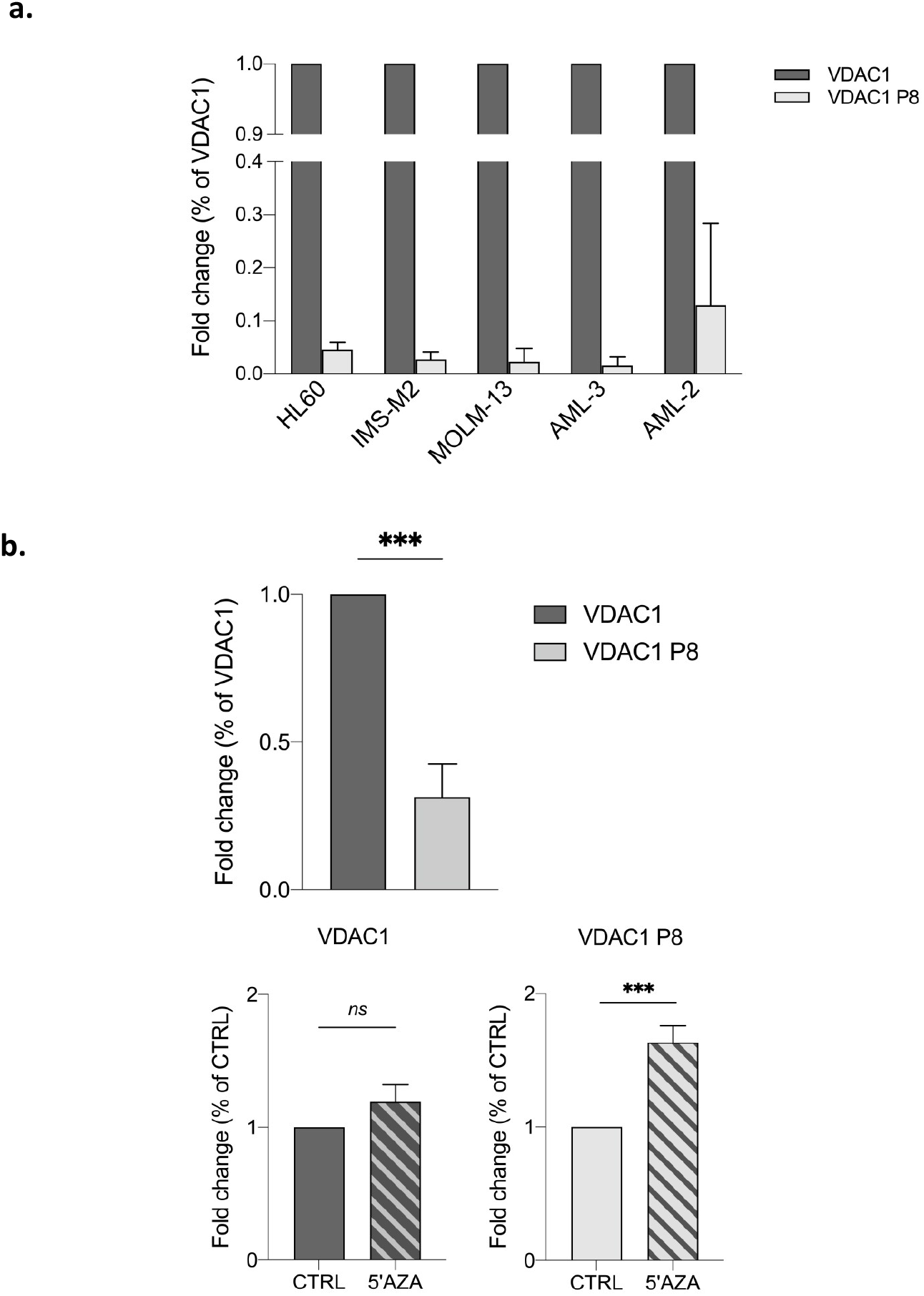
Expression of *VDAC1* gene and *VDAC1P8* pseudogene in HL60 AML cell line. In (a), the expression of *VDAC1* gene and *VDAC1P8* pseudogene was evaluated in different models of leukemia cell line (HL60, IMS-M2, MOLM-13, AML-3, AML-2) by real-time PCR relative quantification. In (b), the expression of *VDAC1* and *VDAC1P8* both assessed in HL60 by real-time PCR relative quantification were also measured following 48h of 5-Aza-2’deoxycytidine treatment. The variation of mRNA in treated samples were expressed relatively to control samples, after normalization with the housekeeping gene *GAPDH*, by the ΔΔCt method. Data were statistically analyzed by T-test and a value of P < 0.05 was considered significant.

Also, the expression level of *VDAC1P8/VDAC1* pair in HL60 was evaluated following treatment with 5-aza-2’-deoxycytidine (AzaD). By inducing demethylation, no change in the transcript level of *VDAC1* was produced while the expression of the *VDAC1P8* pseudogene increased by an average 1.8-fold (Fig. 6b). Probably chromatin methylation, and/or other epigenetic modifications, might modulate the transcriptional changes of *VDAC1* and *VDAC1P8* in AML pathology.

### Modulation of *VDAC1P8* pseudogene expression in *VDAC1* KO leukemic cell lines

We also evaluated the expression of the *VDAC1* gene/ *VDAC1P8* pseudogene pair in the wild-type HAP1 and in the HAP1D*VDAC1* cell lines. HAP1 is a nearly haploid cell line of leukemic origin were by CRISPR/Cas9 editing the *VDAC1* gene has been silenced (*www.horizondiscovery.com*). The HAP1 cells devoid of *VDAC1* were here exploited to analyze, in a leukemic context, the contribution of the *VDAC1* levels to the expression of *VDAC1P8* pseudogene, and vice versa. Interestingly, in HAP1D*VDAC1* cells we found that the expression level of *VDAC1P8* pseudogene, relatively to the parental *VDAC1* gene, is significantly increased in HAP1D*VDAC1* cells compared to HAP1 WT cells (Fig.7a). Indeed, in HAP1 WT cells, where the *VDAC1* transcript is normally expressed, the expression level of the *VDAC1P8* pseudogene is almost 10-fold lower as in above-mentioned cell lines. In contrast, in HAP1D*VDAC1* cells, where *VDAC1* expression is suppressed, the level of the *VDAC1P8* pseudogene is significantly doubled compared to the parental *VDAC1* gene (Fig.7b).

**Figure 7a-b.**
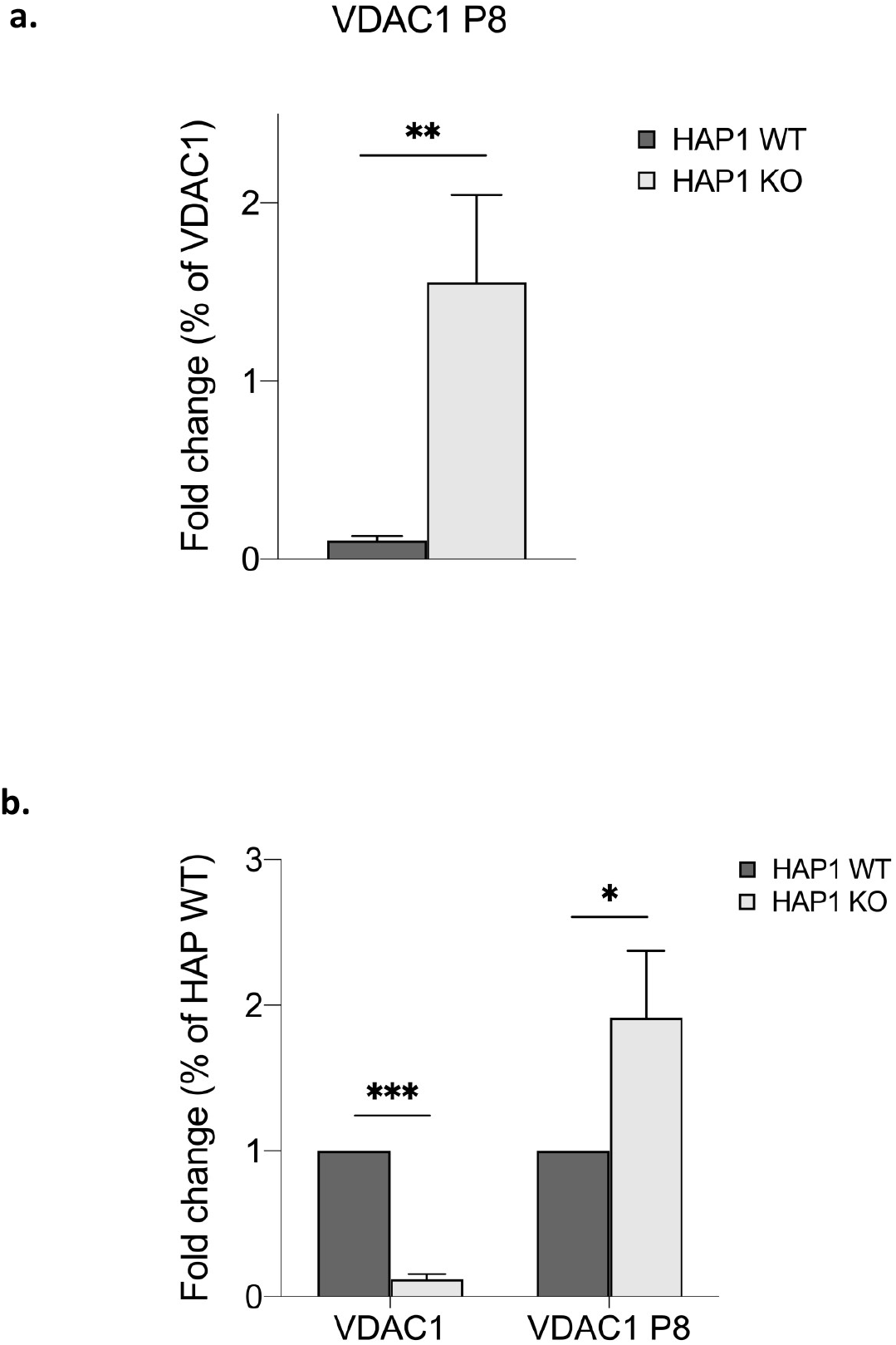
Expression of *VDAC1* gene and *VDAC1P8* pseudogene in *VDAC1* KO leukemic cell lines. The level of *VDAC1* and *VDAC1P8* transcripts expression were evaluated in HAP1 WT and HAP1D*VDAC1* cells by real-time PCR relative quantification. In (a) *VDAC1P8* transcript level was measured relatively to that of *VDAC1*. In (b) the expression level of both *VDAC1* and *VDAC1P8* in HAP1D*VDAC1* cells was measured relatively to their relative amount found in HAP1 WT. After normalization with the housekeeping gene *GAPDH*, by the ΔΔCt method. Data were statistically analyzed by T-test and a value of P < 0.05 was considered significant.

## Discussion

In recent years, transcribed pseudogenes have been increasingly proposed as key regulators of parental gene expression, becoming part of the non-coding RNA network. Since their discovery, almost nothing is known about *VDAC* pseudogenes (26,58). As the factors and circumstances leading to the regulation of *VDAC* genes are still ongoing (5,8,9), understanding whether there is an involvement or a role for their pseudogenes could provide an important key to understanding the whole regulatory network of *VDAC* isoforms expression. This is even more important considering that *VDAC* is already considered a main player in several diseases, and is becoming a prognostic factor and a possible therapeutic target (59,60)

To compensate for the lack of information, in this work we investigated *VDAC* pseudogenes by means of an *in-silico* analysis integrating multi-omic data in terms of genomic features and expression profile, and also validated data in AML cell lines.

We found that among all *VDAC* pseudogenes, *VDAC1P8* appears to reside in an active chromatin region rich in regulatory elements. Exactly, *VDAC1P8* is located between two expressed genes that are controlled by regulatory elements that may direct the expression of the pseudogene transcripts. In this regard, we can speculate that the original *VDAC1* retrotranscription event led to the insertion at the 6q24.2 locus of the exonic sequence that then gave rise to the *VDAC1P8-201* pseudogene. In fact, considering the sequence and the structure of the other 13 annotated alternative *VDAC1P8* transcripts, they do not appear to originate from the *VDAC1* gene and their expression is probably due to the use of different splicing and transcription start sites.

Also, we have analyzed the transcript expression profile of each pseudogene and its parental *VDAC* gene. Expression analysis in healthy tissues, showed ubiquitous activation of some *VDAC1* pseudogenes (*VDAC1P1*, *VDAC1P2*, *VDAC1P6*, *VDAC1P8*), of *VDAC2P1* and of *VDAC3P1*. Intriguingly, the latter is also expressed in testicular germ cells, a tissue in which the cognate gene *VDAC3* plays important functions in spermiogenesis and male fertily (61,62). Some *VDAC1* pseudogenes have been also found to be expressed in cancer, among them some exclusively or predominantly associated with tumor phenotypes. In particular, pseudogenes *VDAC1P1, VDAC1P2, VDAC1P4, VDAC1P11* and *VDAC1P8* correlate with AML. Specifically, the expression profile of *VDAC1P1* and *VDAC1P2* in this analysis largely mirrors that of *VDAC1*, in both healthy and tumor tissues. In contrast, *VDAC1P4, VDAC1P11* and, more interestingly, *VDAC1P8* have an expression trend exactly opposite to that of *VDAC1*. In general, *VDAC1P8* is under-expressed compared to healthy tissue in all tumor samples tested, with the exception of AML where it is highly overexpressed.

These expression data of the VDAC1P8/VDAC1 pair were confirmed in five AML cell lines (myeloblastic, monoblastic, megakaryoblastic) precursors of different blood cells.

The same expression ratio of *VDAC1* and *VDAC1P8* transcripts is also found in the nearly haploid HAP1 wild-type human leukaemic line. In contrast, in the HAP1-*VDAC1* knock-out line, the level of *VDAC1P8* increases significantly, suggesting that the expression of the *VDAC1* gene and its pseudogene *VDAC1P8* may be related to each other.

Also, in regulatory sequences that could control *VDAC1P8* gene we identified 24 putative TFBSs associated with bone marrow, blood and leukemogenesis. Interestingly, common transcriptional factors involved in leukemia are shared in the regulatory sequences of *VDAC1* gene and *VDAC1P8* pseudogene, suggesting that both could play a role in this pathology.

Taken together, these results highlight that *VDAC1P8* is closely associated with leukemia, a finding mentioned in the literature when cancer-specific pseudogenes were identified in chronic lymphocytic leukemia (27).

It is also important to note that previous studies have shown the existence of different DNA methylation rates for pseudogenes and their parent genes (33,63,64).To better understand the significance of *VDAC1* pseudogenes activation patterns in relation to parent gene expression levels, we focused our attention on the DNA methylation status of these pseudogenes and *VDAC1* gene. Exactly, we observed that the methylation levels of the *VDAC* pseudogenes presumably depended on their position within the genome. In particular, the expression of the *VDAC1P8* pseudogene undergoes no or little variation and is presumably under the transcriptional control of adjacent genes, as observed with other pseudogenes (i.e. *LIN28* and *PTEN*) (33,63,64). Indeed, five other genes located at the 6q24.2 locus and controlled by the same regulatory elements acting on *VDAC1P8* correlate with AML.

In addition, we searched in the MethBank database for data on the promoter and gene body methylation profile of the *VDAC1* gene and its pseudogenes, particularly *VDAC1P8*. We found that all *VDAC1*-related pseudogenes exhibit a higher promoter and gene body methylation profile than the *VDAC1* gene, suggesting a normally repressed status but eventually subjected to regulation. Indeed, our experimental data in HL60 show that the regulatory regions of the *VDAC1* gene are essentially undermethylated unlike those of its pseudogene *VDAC1P8*. This result suggests that in AML the pseudogene undergoes a methylation pattern independent of that of the parental gene. In this regard, we could also speculate that chromatin methylation, and epigenetic modifications in general, may contribute to the transcriptional changes of *VDAC1P8 and VDAC1* in AML pathology. Although, the unusual downregulation of the *VDAC1* gene in AML, could be the result of different mechanisms of gene expression regulation. The epigenetic control mechanisms hypothesized here are reflected in other examples found in the literature concerning the transcriptional control of the *VDAC1* gene under extremes conditions. For example, in tumor cells and in placental trophoblasts from patients with recurrent miscarriages, the increase of *EPB41L4A-AS1* lnc-RNA induced the enhancement of *VDAC1* promoter activity affecting histone modification. In this condition, *EPB41L4AAS1* lnc-RNA is considered to regulate and to reprogramme tumor and trophoblast cells metabolism. As a consequence, the activation of oxidative metabolism was triggered by enhanced mitochondrial function (65,66).

In conclusion, our data suggest a role for some VDAC1 pseudogenes in both physiological and pathological processes, particularly in AML. In this context, comparing the expression level of the *VDAC1* gene with the two groups of pseudogenes, those overexpressed and those downregulated, reveals a possible role from ceRNA for these pseudogenes in AML. Considering the inverse relationship between the expression of *VDAC1P4, VDAC1P11* and in particular *VDAC1P8* and *VDAC1*, these pseudogenes in AML could function as competing-endogenous RNA (ceRNA), or rather as a lncRNA able to generate siRNA acting on the transcript of the parental gene, as described in (18). Furthermore, there are numerous examples in the literature of lncRNAs related to leukemia progression (67). With regard to *VDAC1P1, VDAC1P2* these pseudogenes have an expression profile in AML that follows that of the *VDAC1* gene. It is therefore likely that these pseudogenes*-*can act as competitive lncRNA for binding miRNAs or regulatory RNA-binding proteins (RBP) directed to the parental gene, like other pseudogenes (19). This putative mechanism of action correlates very well with the observation that several miRNAs modulating myeloid differentiation (e.g. *miR-20a, miR-106b* and *miR-125b*) (67) or AML (68,69) are also predicted to control *VDAC1* expression (70) using mirSystem, a miRNAtarget prediction software (http://mirsystem.cgm.ntu.edu.tw/index.php) (71). This effect could have important consequences especially considering the role of *VDAC1* protein in apoptosis. Since in leukemia the expression of *VDAC1P8* is considerably higher than in healthy tissue, this pseudogene could be responsible for, or at least contribute to, the down-regulation of *VDAC1* in AML. Given the pro-apoptotic role of VDAC1 protein, in this pathological context a reduced expression could be compatible with the increased cell resistance to apoptosis typical of cancer phenotype. It is also useful to remember that active retrocopies produce lncRNAs, molecules playing many regulatory roles in cancer (72), especially by influencing various cellular mechanisms that promote energy metabolism in cancer (73,74). At the OMM, VDAC serves as a docking site for cytosolic enzymes, such as hexokinase, and is a key protein in mitochondrial apoptosis. Considering the significance of hexokinase for tumor metabolism, its interaction with VDAC and the latter’s role in apoptosis, it is conceivable that lncRNAs modifying the activity of glycolytic enzymes and energy metabolism in cancer have an impact on VDAC function and apoptosis, thus helping to define the tumor phenotype.

## Conclusion

The total number of pseudogenes in the human genome and their function are not yet fully known. It is also unclear why some human genes, such as *VDAC* genes, possess several pseudogenes, differently localized from their parental genes, while others possess only one or a few, and still others none. Our search for orthologous genes indicated strong conservation throughout primate evolution of all *VDAC1* pseudogenes. This leads us to consider *VDAC1* pseudogenes advantageous to primates and to hypothesize that there was selective pressure towards their maintenance already at birth in ancestral primates. This is not surprising considering that *VDAC1* is a very important housekeeping gene under stress conditions, when mitochondrial function must be ensured in cells (75).

Overall, this is the first comprehensive survey of human *VDAC* pseudogenes reported in the literature, and the results presented here provide important insights into their relevance to human genome expression. We found evidence for their activation and importance as human-specific regulatory elements. In particular, *VDAC1P8* appears to be particularly involved in hematopoiesis and emerges as a putative AML-specific diagnostic and prognostic factor. This leads to the evaluation of its regulatory mechanism as a possible target in the fight against the disease. The low survival rate of AML patients ignites the demand for new therapies for this widespread malignancy (84). More in-depth studies are therefore desirable to elucidate the mechanism by which *VDAC1P8* (and other *VDAC* pseudogenes) influence the expression of parental genes in AML, i.e. how their ncRNAs act on regulatory RNA, DNA or RBPs. So far, we have provided data supporting the association of the *VDAC1P8* pseudogene with AML as well as a suitable detection method to distinguish it from its parental gene *VDAC1*. This lays the foundation for its possible use as a biomarker for AML.

## Methods

### Data Collection Process

Specific repositories used to retrieve data on *VDAC* pseudogenes were Ensemble (https://www.ensembl.org/index.html), NCBI Entrez Gene (https://www.ncbi.nlm.nih.gov/), UCSC Genomic Browser (https://genome.ucsc.edu), Hoppsigen database (http://pbil.univ-lyon1.fr/databases/hoppsigen.html) (76), pseudogene.org. by Yale Gerstein Group (77), DreamBase (https://rna.sysu.edu.cn/dreamBase/) (78) and PseudoFun (https://integrativeomics.shinyapps.io/pseudofun_app/) (79).

### Genomic data analysis

UCSC Genomic Browser (assembly GRCh38/hg38) was used to analyze the genomic structure of *VDAC* pseudogenes and gene regulatory data from hub tracks’ selection. This included Pseudogene Annotation Set of GENCODE v.38lift37 (Ensemble 104), Eukaryotic Promoter Database (EPD) v.4-6, CpG islands track, Genotype-Tissue Expression (GTEx) RNA-seq v.8 (2019), ChIP-Seq data for RNA polymerase II, H3K4me3 and H3K4me1, and chromatin state segmentation by Hidden Markov Model from ENCODE/Broad project. GRCh37/hg19 assembly was used because it provided a more suitable representation of the genomic trait for data visualization and interpretation. The range of analysis of the genomic context of each pseudogene was set about 1000 kb upstream and downstream from the Refseq annotation. In addition, comparative genomics data were acquired from the Ensembl vertebrate genome browser (https://www.ensembl.org/index.html) to synteny search between human *VDAC* genes or their pseudogenes and more evolved primates. Also, the GeneCards - The Human Gene Database (https://www.genecards.org/) was queried to refine the genomic information related to *VDAC* pseudogenes.

### Transcriptomic data analysis

GEPIA (Gene Expression Profiling Interactive Analysis) web-based tool and GEPIA2 version were accessed in order to collect gene expression profiles of *VDAC* pseudogenes and their parental genes, in both tumor and healthy tissues (80,81). GEPIA provides data of TCGA (82) and GTEx (Genotype-Tissue Expression project) dataset release v.8 (https://www.gtexportal.org/home/). Expression Atlas (www.ebi.ac.uk/gxa/home) (83) was also exploited for the analysis of adult and fetal human tissues in RIKEN functional annotation of the mammalian genome 5 (FANTOM5) project (https://fantom.gsc.riken.jp/5/) (84). In addition, GWAS Catalog at https://www.ebi.ac.uk/gwas/home was queried for gene-linked phenotypes/functions in genome-wide association studies.

### Sequence Analysis

Sequence alignment analysis was running at free software online (http://multalin.toulouse.inra.fr/multalin/). The percentage of homology between the examined sequences was obtained by BLAST website (https://blast.ncbi.nlm.nih.gov/Blast.cgi). The search for orthologous genes was performed using the BLAST/BAT tool at Ensembl.

### Prediction analysis of transcription factor binding sites (TFBSs) in the *VDAC* pseudogenes regulatory regions

The prediction of potential transcription factors (TFs) and their transcription factor binding sites (TFBSs) on sequences associated with human *VDAC1* pseudogenes was conducted by hTFtarget database (http://bioinfo.life.hust.edu.cn/hTFtarget#!/) (85). Each record of candidate TFs for query gene provides the chromosomal location of chromatin immunoprecipitation sequencing (ChIP-seq) peaks in different experimental conditions or cell lines. The association between candidate TF and target sequences is computed through a machine learning method and accumulated data of ChIP-seq peaks, epigenetic modification status of TFBSs and targets, co-regulation of TFs of the same targets, and co-association of TFs.

### DNA methylation analysis

DNA methylation profiles of both regulatory sequences and gene bodies of *VDAC1* and *VDAC1P8* were investigated using data of single-base resolution methylomes (SRMs) retrieved from MethBank v.4.1 (Sep 2020) (https://ngdc.cncb.ac.cn/methbank) (86). Since MethBank contains a large amount of SRM replicates both from healthy and pathological condition (prevalently lung and prostate carcinomas), our search strategy filtered SRM data for ‘human healthy tissues’ selecting epigenetic change of guanine-cytosine (‘GC’) base pair found in the ‘promoter’ and ‘gene body’. The average methylation data are calculated as β-value that reflects the methylation intensity at each CpG site. β-values of 0–1 (represented from 0 to 1) indicate signifying percent methylation, from 0 to 100%, respectively, for each CpG site. For illustration purpose, in case of some tissue replicates (adipose, adrenal gland, bladder, breast, lung, placenta, prostate, psoas) we averaged their β-values showing a single record for each sample. We also averaged samples belonging to the same apparatus clustering them into following systems: circulatory (aorta, carotid, right atrium, right and left ventricles, whole blood); digestive (pancreas, small and large intestine, stomach, colon, sigmoid colon, esophagus, esophagus squamous epithelium, adjacent esophageal tissue, gastric); lymphatic (spleen, thymus), nervous (anterior cingulate cortex, cerebral cortex (fetal), frontal lobus, hippocampus, nucleus accumbens, prefrontal cortex (Broadman area 9), retina).

### Cell cultures and treatment

The acute myeloid leukemia model cell lines (OCI AML2, OCI AML3, MOLM13, IMS M2) from American Type Culture Collection (Manassas, VA, USA) and HL60 (from DSMZ cell cultures, Braunschweig, German) were cultured in RPMI-1640 medium (Sigma Chemical Co. - St. Louis, MO, USA) supplemented with 10% fetal bovine serum, 2 mM L-Glutamine, 100 U/ml Penicillin and 100 μg/ml Streptomycin (EuroClone, Pero, MI, Italy). The cells were grown at 37°C in a humidified atmosphere containing 5% CO2 and dissociated with 0,25% trypsin-0,02% EDTA solution. The HAP1 parental (C631) and knock-out for VDAC1 (HZGHC005706c002) cell lines were purchased from Horizon Discovery (Waterbeach, UK).

Cells were maintained as in (87), as was the monitoring of their proliferation. For cell treatments, 5-Aza-2’deoxycytidine (AzaD) was purchased from Sigma Chemical Co. (St. Louis, MO, USA) and resuspended in dimethyl sulfoxide (DMSO) at the concentration of 50 mg/mL, respectively. The cells were seeded at 0,1 x10^6^/ml in 25 cm^2^ culture flasks (NUNC, Thermo Scientific, Roskilde, Denmark) and treated with 5 μM AzaD for 48 hours.

### Real-time amplification

Real-time amplification was performed in a Mastercycler EP Realplex (Eppendorf) in 96-well plates. cDNAs from leukemia cells line HL60 were used as template for the experiments. The reaction mixture was composed by 1 μl cDNA, 0.2 μM gene specific primers pairs for *VDAC1, VDAC1P8-201* and *GAPDH* (Suppl. Table 5) and 12.5 μl of master mix (QuantiFast SYBR Green PCR kit, Qiagen). Analysis of relative expression level was performed using the housekeeping *GAPDH* gene as internal calibrator by the ΔΔCt method. Three independent experiments were performed. Data were statistically analyzed by Student t-test. Values of * p<0.05, ** p<0.01 and *** p<0.001 were taken as significant.

## Supporting information

Supplementary Informations

## List of abbreviations

AzaD: 5-aza-2’-deoxycytidine;
AML: acute myeloid leukemia;
ceRNA: ChlP-seq, chromatin immunoprecipitation sequencing;
circRNA: circular RNA;
competitive endogenous RNA;
EPD: Eukaryotic Promoter Database;
GEPIA: Gene Expression Profiling Interactive Analysis;
GTEX: Genotype-Tissue Expression;
HK: hexokinase;
lncRNA: long non-coding RNA;
miRNA: microRNA;
OMM: outer mitochondrial membrane;
TF: transcription factor;
TFBS: transcription factor binding site;
siRNA: short interfering RNAs;
RBP: RNA-binding proteins;
SRMs: single-base resolution methylomes;
TPM: transcript per million;
VDACs: Voltage dependent anion selective channels.

## Acknowledgments

The authors are grateful to Dr. Daniele Tibullo (University of Catania) for the gift of the AML cell lines used in the work. Authors also acknowledge Fondi di Ateneo 2020–2022 and Linea Open Access, University di Catania.

## Author Contributions

XGP developed the pipelines, did most of the bioinformatic analyses, and wrote together PR the initial drafts of the manuscript. PR performed also some bioinformatics analysis and analysed the data. FG performed the sequence alignments. FZ and FG performed the analysis to search for TFs. AO and DL performed all cell experiments.

AM conceived the study, performed some bioinformatics analysis and wrote the final drafts of the manuscript. VDP and FB revised and edited the text. XGP, FG, AM contributed to the final version of the manuscript. All authors read and approved the final manuscript.

## Data availability

All data generated or analyzed during this study are included in this published article.

## Declarations

### Ethics approval and consent to participate

Not applicable

### Consent for publication

Not applicable

