## Supplementary Informations for "Human *VDAC* pseudogenes: an emerging role for *VDAC1P8* pseudogene in acute myeloid leukemia"

### Supplementary Figures

Suppl. Fig. 1a-b

**a**

| ***VDAC1P8* transcripts in Ensembl database (release 105)** | | | | |
|---|---|---|---|---|
| **Transcript ID** | **Name** | **bp** | **Biotype** | **Exon n°** |
| ENST00000610068.5 | *VDAC1P8-210* | 1251 | Processed transcript | 5 |
| ENST00000589489.1 | *VDAC1P8-204* | 997 | Retained intron | 2 |
| ENST00000438118.6 | *VDAC1P8-203* | 987 | Retained intron | 3 |
| ENST00000619849.4 | *VDAC1P8-213* | 982 | Retained intron | 4 |
| ENST00000622321.1 | *VDAC1P8-214* | 902 | Retained intron | 3 |
| ENST00000612298.1 | *VDAC1P8-212* | 886 | Retained intron | 2 |
| ENST00000406025.2 | *VDAC1P8-201* | 853 | Transcribed processed pseudogene | 1 |
| ENST00000593045.5 | *VDAC1P8-208* | 780 | Retained intron | 3 |
| ENST00000589563.5 | *VDAC1P8-205* | 752 | Processed transcript | 4 |
| ENST00000590703.5 | *VDAC1P8-206* | 752 | Retained intron | 2 |
| ENST00000415586.5 | *VDAC1P8-202* | 702 | Processed transcript | 3 |
| ENST00000593175.1 | *VDAC1P8-209* | 698 | Retained intron | 3 |
| ENST00000591189.5 | *VDAC1P8-207* | 640 | Processed transcript | 5 |
| ENST00000611810.1 | *VDAC1P8-211* | 422 | Processed transcript | 3 |

**b**

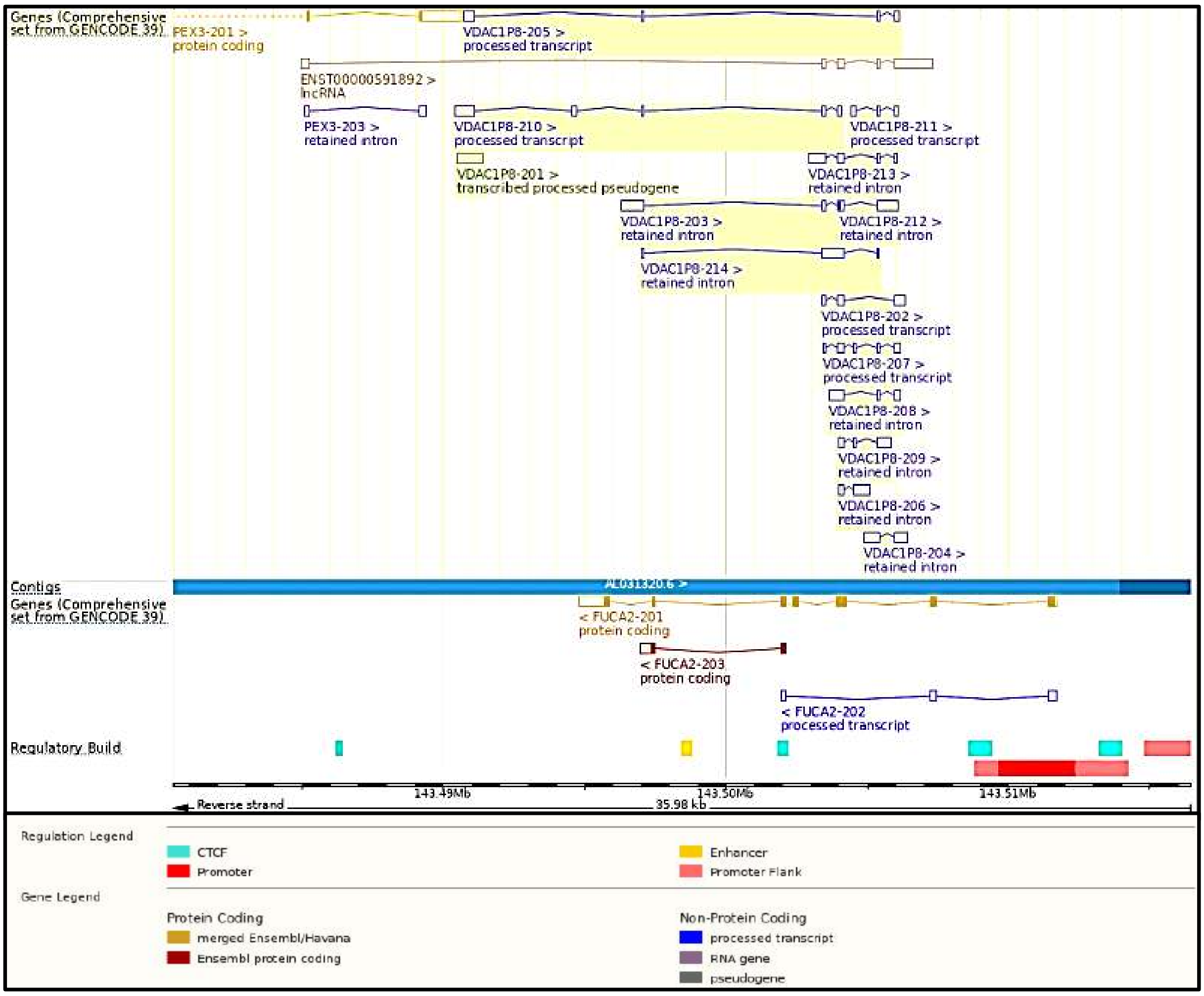

**Suppl. Fig. 1a-b. Summary of annotated transcripts for the *VDAC1P8* pseudogene from GENCODE v.39 Ensembl 105.** In (a) the table reports gene annotations of 14 transcripts for *VDAC1P8* reported on Ensembl database release 105. In (b) the screenshot modified by GENCODE v.39 illustrates some main structural and regulatory features of the variants' alignment.

**Suppl. Fig. 2**

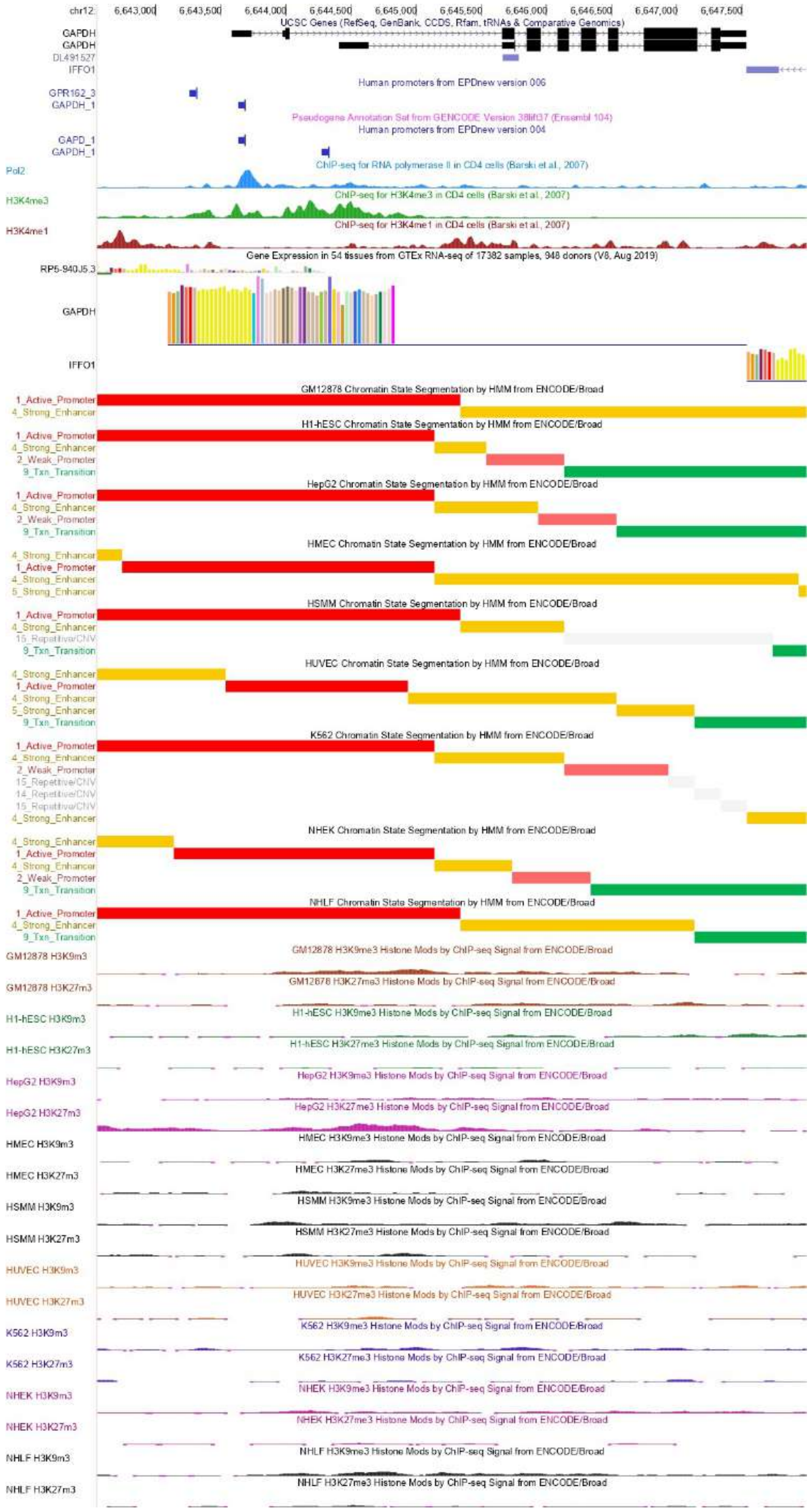
chr12:
6,643,000
6,643,500
6,644,000
6,644,500
6,645,000
6,645,500
6,646,000
6,646,500
6,647,000
6,647,500
UCSC Genes (RefSeq, GenBank, CCDS, Rfam, tRNAs & Comparative Genomics)
GAPDH
GAPDH
DL491527
IFFO1
Human promoters from EPDnew version 006
GPR162_3
GAPDH_1
Pseudogene Annotation Set from GENCODE Version 38lift37 (Ensembl 104)
Human promoters from EPDnew version 004
GAPD_1
GAPDH_1
ChIP-seq for RNA polymerase II in CD4 cells (Barski et al., 2007)
Pol2
H3K4me3
ChIP-seq for H3K4me3 in CD4 cells (Barski et al., 2007)
H3K4me1
ChIP-seq for H3K4me1 in CD4 cells (Barski et al., 2007)
Gene Expression in 54 tissues from GTEx RNA-seq of 17382 samples, 948 donors (V8, Aug 2019)
RP5-940J5.3
GAPDH
IFFO1
GM12878 Chromatin State Segmentation by HMM from ENCODE/Broad
H1-hESC Chromatin State Segmentation by HMM from ENCODE/Broad
HepG2 Chromatin State Segmentation by HMM from ENCODE/Broad
HMEC Chromatin State Segmentation by HMM from ENCODE/Broad
HSMM Chromatin State Segmentation by HMM from ENCODE/Broad
HUVEC Chromatin State Segmentation by HMM from ENCODE/Broad
K562 Chromatin State Segmentation by HMM from ENCODE/Broad
NHEK Chromatin State Segmentation by HMM from ENCODE/Broad
NHLF Chromatin State Segmentation by HMM from ENCODE/Broad
1_Active_Promoter
2_Weak_Promoter
4_Strong_Enhancer
5_Strong_Enhancer
9_Txn_Transition
14_Repetitive/CNV
15_Repetitive/CNV
GM12878 H3K9m3
GM12878 H3K9me3 Histone Mods by ChIP-seq Signal from ENCODE/Broad
GM12878 H3K27m3
GM12878 H3K27me3 Histone Mods by ChIP-seq Signal from ENCODE/Broad
H1-hESC H3K9m3
H1-hESC H3K9me3 Histone Mods by ChIP-seq Signal from ENCODE/Broad
H1-hESC H3K27m3
H1-hESC H3K27me3 Histone Mods by ChIP-seq Signal from ENCODE/Broad
HepG2 H3K9m3
HepG2 H3K9me3 Histone Mods by ChIP-seq Signal from ENCODE/Broad
HepG2 H3K27m3
HepG2 H3K27me3 Histone Mods by ChIP-seq Signal from ENCODE/Broad
HMEC H3K9m3
HMEC H3K9me3 Histone Mods by ChIP-seq Signal from ENCODE/Broad
HMEC H3K27m3
HMEC H3K27me3 Histone Mods by ChIP-seq Signal from ENCODE/Broad
HSMM H3K9m3
HSMM H3K9me3 Histone Mods by ChIP-seq Signal from ENCODE/Broad
HSMM H3K27m3
HSMM H3K27me3 Histone Mods by ChIP-seq Signal from ENCODE/Broad
HUVEC H3K9m3
HUVEC H3K9me3 Histone Mods by ChIP-seq Signal from ENCODE/Broad
HUVEC H3K27m3
HUVEC H3K27me3 Histone Mods by ChIP-seq Signal from ENCODE/Broad
K562 H3K9m3
K562 H3K9me3 Histone Mods by ChIP-seq Signal from ENCODE/Broad
K562 H3K27m3
K562 H3K27me3 Histone Mods by ChIP-seq Signal from ENCODE/Broad
NHEK H3K9m3
NHEK H3K9me3 Histone Mods by ChIP-seq Signal from ENCODE/Broad
NHEK H3K27m3
NHEK H3K27me3 Histone Mods by ChIP-seq Signal from ENCODE/Broad
NHLF H3K9m3
NHLF H3K9me3 Histone Mods by ChIP-seq Signal from ENCODE/Broad
NHLF H3K27m3
NHLF H3K27me3 Histone Mods by ChIP-seq Signal from ENCODE/Broad

**Suppl. Fig. 2. Chromatin state and genomic features of *GAPDH* gene from UCSC Genome Browser GRCh37/hg19.** The genomic context of *GAPDH* set around 1000 Kb upstream and downstream of the annotated Refseq is shown. The selected regulatory hub tracks are Pseudogene Annotation Set from GENCODE v.38lift37 Ensemble 104, Eukaryotic Promoter Database EPD v.4-6, CpG island track, Genotype-Tissue Expression GTEx RNA-seq v.8 2019, ChIP-Seq data for RNA polymerase II, H3K4me3 and H3K4me1, used as markers of transcriptional activation, while H3K9me3 and H3K27me3 are markers of transcriptional repression, and chromatin state segmentation by Hidden Markov Model from the ENCODE/Broad project of nine different cell lines (GM12878, H1-hESC, HepG2, HMEC, HUVEC, K562, NHEK, NHLF) identified using the following different colors: bright red=active promoter; light red= week promoter; orange= strong enhancer; dark green= transcriptional transition/elongation.

**Suppl. Fig. 3**

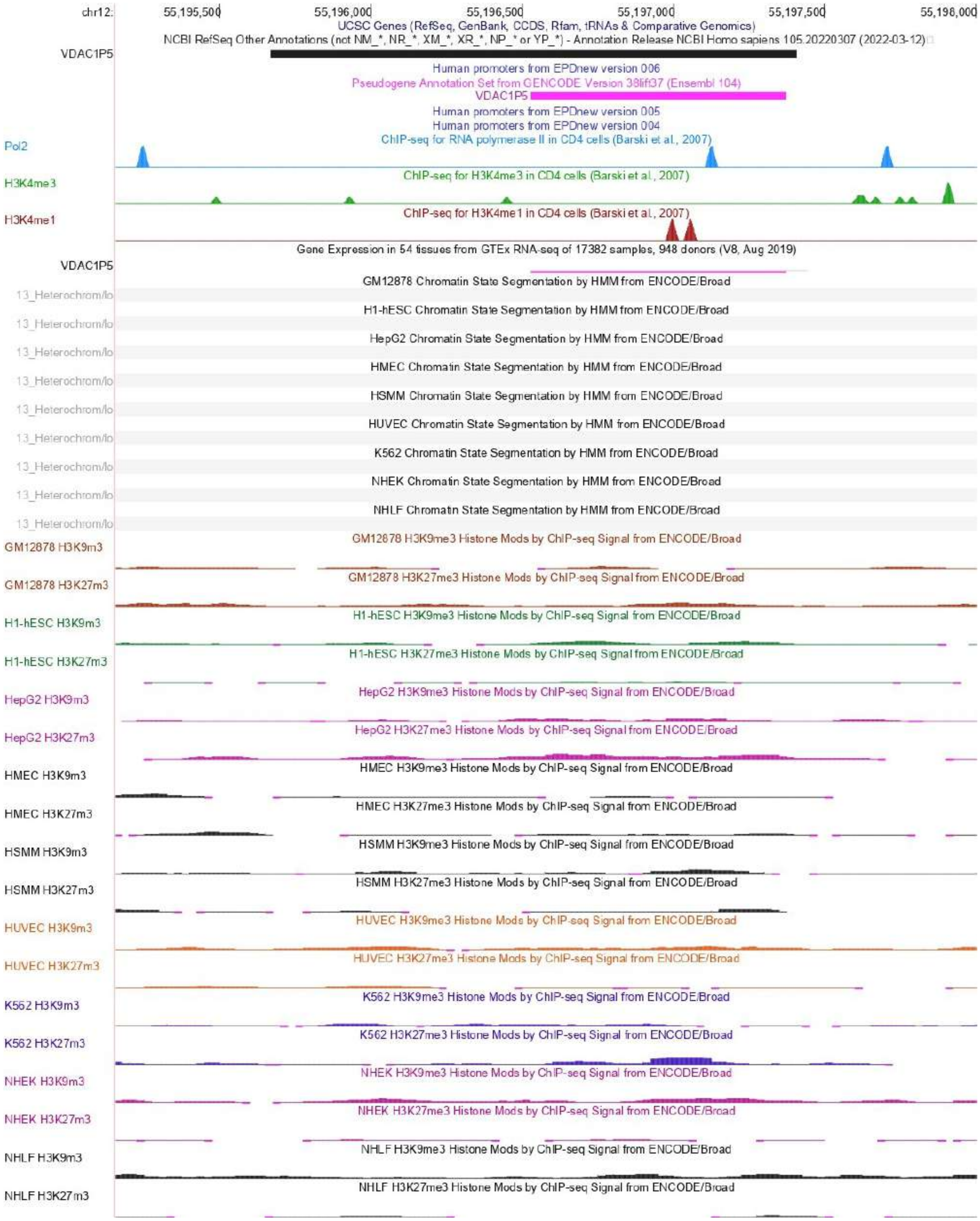
chr12:
55,195,500
55,196,000
55,196,500
55,197,000
55,197,500
55,198,000
UCSC Genes (RefSeq, GenBank, CCDS, Rfam, tRNAs & Comparative Genomics)
NCBI RefSeq Other Annotations (not NM_*, NR_*, XM_*, XR_*, NP_* or YP_*) - Annotation Release NCBI Homo sapiens 105.20220307 (2022-03-12)
VDAC1P5
Human promoters from EPDnew version 006
Pseudogene Annotation Set from GENCODE Version 38lift37 (Ensembl 104)
VDAC1P5
Human promoters from EPDnew version 005
Human promoters from EPDnew version 004
Pol2
ChIP-seq for RNA polymerase II in CD4 cells (Barski et al., 2007)
H3K4me3
ChIP-seq for H3K4me3 in CD4 cells (Barski et al., 2007)
H3K4me1
ChIP-seq for H3K4me1 in CD4 cells (Barski et al., 2007)
Gene Expression in 54 tissues from GTEx RNA-seq of 17382 samples, 948 donors (V8, Aug 2019)
VDAC1P5
GM12878 Chromatin State Segmentation by HMM from ENCODE/Broad
13_Heterochrom/lo
H1-hESC Chromatin State Segmentation by HMM from ENCODE/Broad
13_Heterochrom/lo
HepG2 Chromatin State Segmentation by HMM from ENCODE/Broad
13_Heterochrom/lo
HMEC Chromatin State Segmentation by HMM from ENCODE/Broad
13_Heterochrom/lo
HSMM Chromatin State Segmentation by HMM from ENCODE/Broad
13_Heterochrom/lo
HUVEC Chromatin State Segmentation by HMM from ENCODE/Broad
13_Heterochrom/lo
K562 Chromatin State Segmentation by HMM from ENCODE/Broad
13_Heterochrom/lo
NHEK Chromatin State Segmentation by HMM from ENCODE/Broad
13_Heterochrom/lo
NHLF Chromatin State Segmentation by HMM from ENCODE/Broad
13_Heterochrom/lo
GM12878 H3K9m3
GM12878 H3K9me3 Histone Mods by ChIP-seq Signal from ENCODE/Broad
GM12878 H3K27m3
GM12878 H3K27me3 Histone Mods by ChIP-seq Signal from ENCODE/Broad
H1-hESC H3K9m3
H1-hESC H3K9me3 Histone Mods by ChIP-seq Signal from ENCODE/Broad
H1-hESC H3K27m3
H1-hESC H3K27me3 Histone Mods by ChIP-seq Signal from ENCODE/Broad
HepG2 H3K9m3
HepG2 H3K9me3 Histone Mods by ChIP-seq Signal from ENCODE/Broad
HepG2 H3K27m3
HepG2 H3K27me3 Histone Mods by ChIP-seq Signal from ENCODE/Broad
HMEC H3K9m3
HMEC H3K9me3 Histone Mods by ChIP-seq Signal from ENCODE/Broad
HMEC H3K27m3
HMEC H3K27me3 Histone Mods by ChIP-seq Signal from ENCODE/Broad
HSMM H3K9m3
HSMM H3K9me3 Histone Mods by ChIP-seq Signal from ENCODE/Broad
HSMM H3K27m3
HSMM H3K27me3 Histone Mods by ChIP-seq Signal from ENCODE/Broad
HUVEC H3K9m3
HUVEC H3K9me3 Histone Mods by ChIP-seq Signal from ENCODE/Broad
HUVEC H3K27m3
HUVEC H3K27me3 Histone Mods by ChIP-seq Signal from ENCODE/Broad
K562 H3K9m3
K562 H3K9me3 Histone Mods by ChIP-seq Signal from ENCODE/Broad
K562 H3K27m3
K562 H3K27me3 Histone Mods by ChIP-seq Signal from ENCODE/Broad
NHEK H3K9m3
NHEK H3K9me3 Histone Mods by ChIP-seq Signal from ENCODE/Broad
NHEK H3K27m3
NHEK H3K27me3 Histone Mods by ChIP-seq Signal from ENCODE/Broad
NHLF H3K9m3
NHLF H3K9me3 Histone Mods by ChIP-seq Signal from ENCODE/Broad
NHLF H3K27m3
NHLF H3K27me3 Histone Mods by ChIP-seq Signal from ENCODE/Broad

**Suppl. Fig. 3. Chromatin state and genomic features of *VDAC1P5* gene from UCSC Genome Browser GRCh37/hg19.** The genomic context of *VDAC1P5* set around 1000 Kb upstream and downstream of the annotated Refseq is shown. The selected regulatory hub tracks are Pseudogene Annotation Set from GENCODE v.38lift37 Ensemble 104, Eukaryotic Promoter Database EPD v.4-6, CpG island track, Genotype-Tissue Expression GTEx RNA-seq v.8 2019, ChIP-Seq data for RNA polymerase II, H3K4me3 and H3K4me1, used as markers of transcriptional activation, H3K9me3 and H3K27me3 are markers of transcriptional repression, and chromatin state segmentation by Hidden Markov Model from the ENCODE/Broad project of nine different cell lines (GM12878, H1-hESC, HepG2, HMEC, HUVEC, K562, NHEK, NHLF) colored in grey to indicate the heterochromatin state.

**Suppl. Fig. 4a-b**

**a**

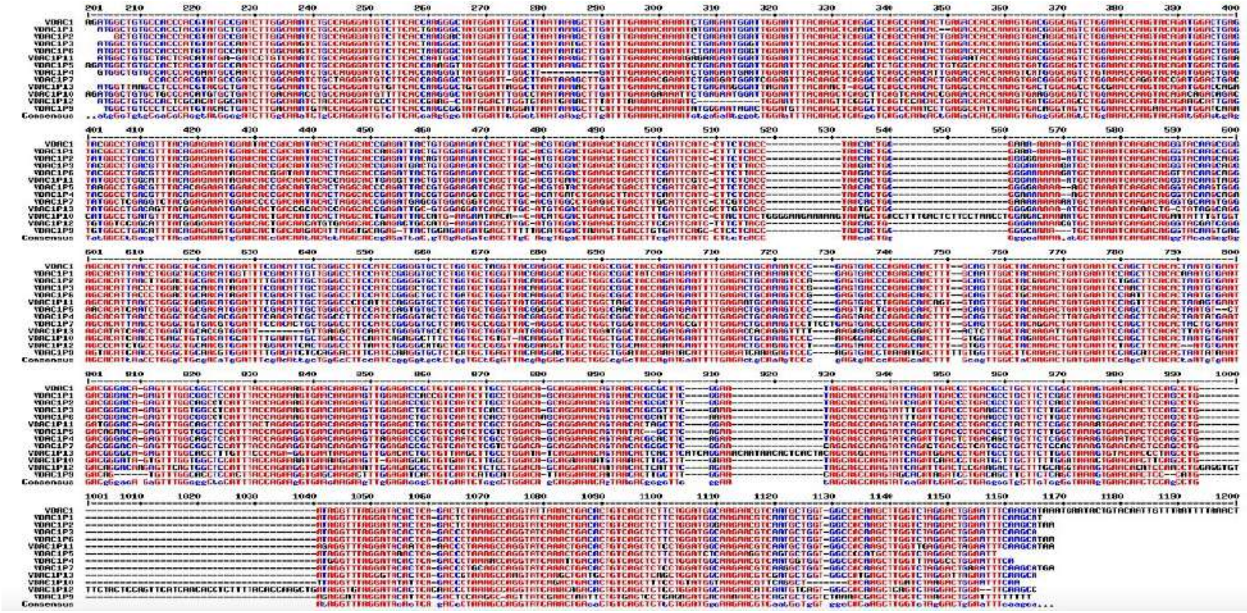

**b**

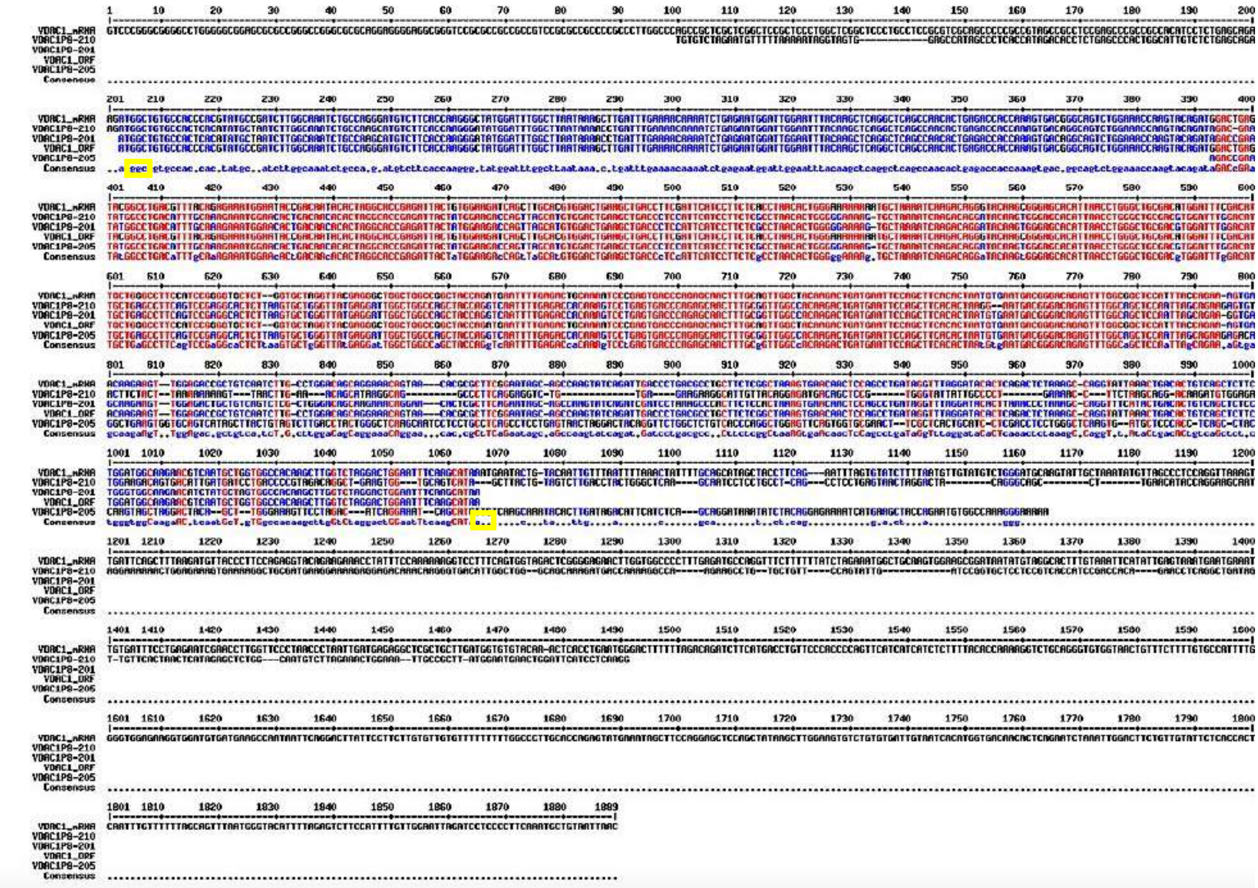

**Suppl. Fig. 4a-b. Sequence multi-alignments among *VDAC1* and pseudogenes.** In (a) the multi-alignment encompasses *VDAC1* against *VDAC1P1-7* and *VDAC1P9-13*. In this picture, the *VDAC1* mRNA sequence is shown starting at nucleotide 201 because its start ATG is located at 203-205 position and also because the previous sequence stretches nucleotides 1-200 does not align with the *VDAC1* pseudogenes sequences. In (b), the multi-alignment is among *VDAC1* and splicing variants of *VDAC1P8 VDAC1P8-201, -205* and -*210*. The start and end codons of the *VDAC1* coding sequence are boxed in yellow.

**Suppl. Fig. 5**

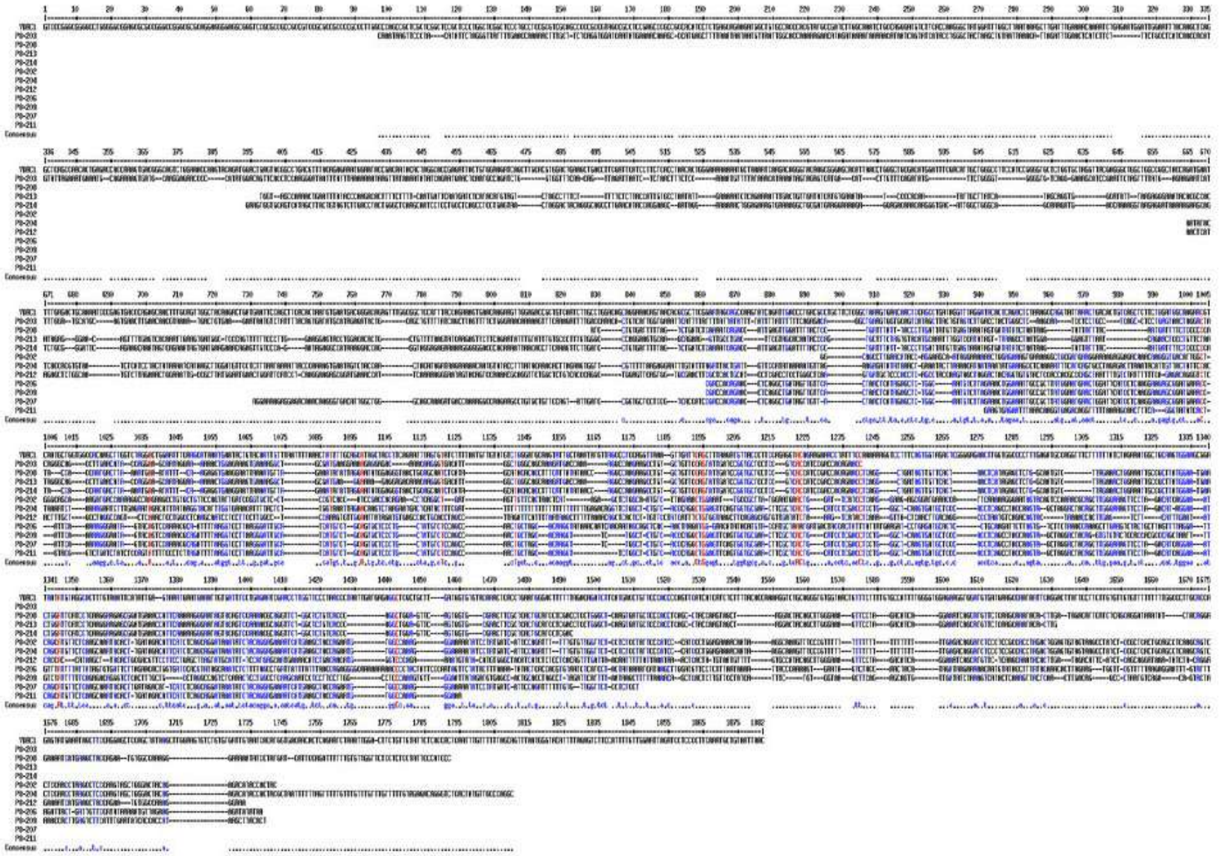

**Suppl. Fig. 5. Multi-alignment between the *VDAC1* mRNA and some *VDAC1P8* alternative transcripts sequences.** Alignment achieved by http://multalin.toulouse.inra.fr/multalin/ setting the default parameters. The regions with high consensus value (90% setting) are shown in red, while regions with low consensus value (50% setting) are shown in blue.

**Suppl. Fig. 6a-b**

**a.** **Average methylation of *VDAC1, VDAC1P2, VDAC1P4, VDAC1P8 and VDAC1P11* promoter**

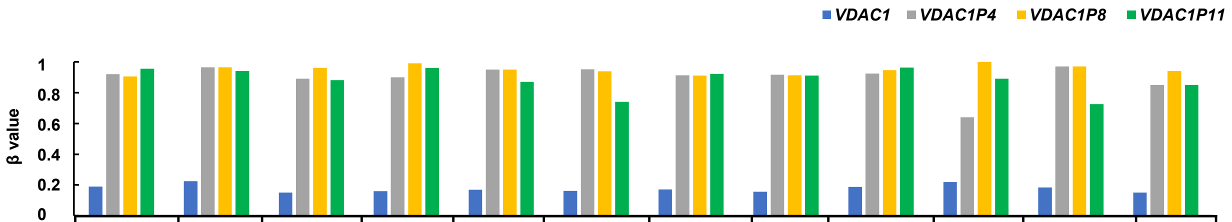


| | Adipose | Adrenal gland | Bladder | Breast | Central nervous system | Circulatory system | Digestive system | Lung | Lymphatic system | Placenta | Prostate | Psoas |
|---|---|---|---|---|---|---|---|---|---|---|---|---|
| ***VDAC1*** | 0.19 | 0.23 | 0.15 | 0.16 | 0.17 | 0.16 | 0.17 | 0.16 | 0.19 | 0.22 | 0.19 | 0.15 |
| ***VDAC1P4*** | 0.92 | 0.97 | 0.89 | 0.9 | 0.95 | 0.95 | 0.91 | 0.92 | 0.92 | 0.64 | 0.97 | 0.85 |
| ***VDAC1P8*** | 0.91 | 0.97 | 0.96 | 0.99 | 0.95 | 0.94 | 0.91 | 0.91 | 0.95 | 1 | 0.97 | 0.94 |
| ***VDAC1P11*** | 0.96 | 0.94 | 0.88 | 0.96 | 0.87 | 0.74 | 0.92 | 0.91 | 0.96 | 0.89 | 0.73 | 0.85 |

**b.** **Average methylation of *VDAC1, VDAC1P2, VDAC1P4, VDAC1P8 and VDAC1P11* gene body**

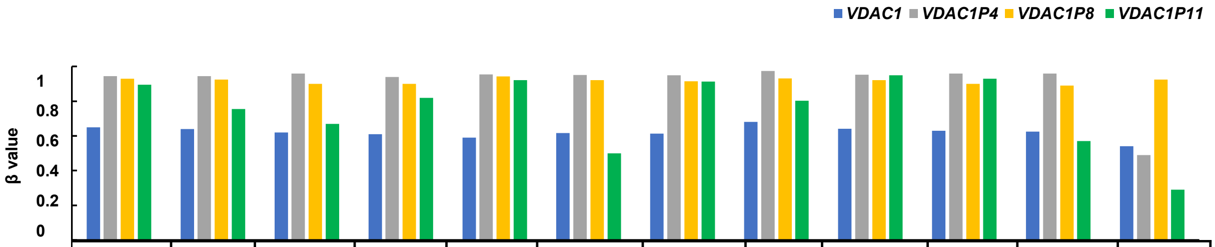


| | Adipose | Adrenal gland | Bladder | Breast | Central nervous system | Circulatory system | Digestive system | Lung | Lymphatic system | Placenta | Prostate | Psoas |
|---|---|---|---|---|---|---|---|---|---|---|---|---|
| ***VDAC1*** | 0.65 | 0.64 | 0.62 | 0.61 | 0.59 | 0.62 | 0.61 | 0.68 | 0.64 | 0.63 | 0.63 | 0.54 |
| ***VDAC1P4*** | 0.95 | 0.95 | 0.96 | 0.94 | 0.95 | 0.95 | 0.95 | 0.97 | 0.95 | 0.96 | 0.96 | 0.49 |
| ***VDAC1P8*** | 0.93 | 0.93 | 0.9 | 0.9 | 0.94 | 0.92 | 0.92 | 0.93 | 0.92 | 0.9 | 0.89 | 0.93 |
| ***VDAC1P11*** | 0.9 | 0.76 | 0.67 | 0.82 | 0.92 | 0.5 | 0.91 | 0.8 | 0.95 | 0.93 | 0.57 | 0.29 |

**Suppl. Fig. 6a-b. Methylation levels of *VDAC1P1, VDAC1P4, VDAC1P8* and *VDAC1P11* gene in putative promoter (a) sequence and gene body (b).** **The average methylation of* different healthy human samples from single-based resolution methylomes (SRMs) are provided by MethBank v.4.1 (https://ngdc.cncb.ac.cn/methbank). *SRMs data are calculated as β-Value that reflects the methylation intensity at each CpG site. β-Values of 0–1 (represented from 0 to 1) indicate signifying percent methylation, from 0 to 100%, respectively, for each CpG site.*

### Supplementary Tables

**Suppl. Table 1. Expression profiles of *VDAC1-3* genes from GEPIA server across 31 tumor and paired normal tissues.** The median expression of Log2TPM+1 of certain tumor type or normal tissue based on TCGA and GTEX respectively. TPM= transcript per million.

**color gradient**

**Lowest value** — **Highest value**

| Tumor | Acronym | *VDAC1* Tumor | *VDAC1* Normal | *VDAC2* Tumor | *VDAC2* Normal | *VDAC3* Tumor | *VDAC3* Normal |
|---|---|---|---|---|---|---|---|
| adrenocortical cancer | **ACC** | 207,46 | 104,87 | 144,85 | 133,77 | 77,43 | 78,29 |
| bladder urothelial carcinoma | **BLCA** | 173,78 | 90,31 | 125,35 | 218,22 | 86,08 | 92,84 |
| breast invasive carcinoma | **BRCA** | 177,33 | 86,76 | 154,7 | 127,06 | 108,89 | 70,92 |
| cervical & endocervical cancer | **CESC** | 158,2 | 64,73 | 190,27 | 137,28 | 103,83 | 72,91 |
| cholangiocarcinoma | **CHOL** | 284,7 | 81,63 | 143,78 | 35,87 | 82,75 | 30,96 |
| colon adenocarcinoma | **COAD** | 296,71 | 87,18 | 186,02 | 178,23 | 118,34 | 102,25 |
| diffuse large B-cell lymphoma | **DLBC** | 148,49 | 11,09 | 115,67 | 21,44 | 68,4 | 16,27 |
| esophageal carcinoma | **ESCA** | 182,57 | 134,13 | 199,48 | 260 | 79,45 | 78,87 |
| glioblastoma multiforme | **GBM** | 174,33 | 135,48 | 117,49 | 99,21 | 118,25 | 110,12 |
| head & neck squamous cell carcinoma | **HNSC** | 174,36 | 169,97 | 231,46 | 230,84 | 81,78 | 87,03 |
| kidney chromophobe | **KICH** | 280,04 | 158,22 | 138,88 | 145,97 | 92,28 | 87,15 |
| kidney clear cell carcinoma | **KIRC** | 233,49 | 187,22 | 161,16 | 169,61 | 87,87 | 91,55 |
| kidney papillary cell carcinoma | **KIRP** | 173,63 | 172,31 | 120,69 | 142,9 | 90,04 | 83,79 |
| acute myeloid leukemia | **LAML** | 80,01 | 221,12 | 113,54 | 446,32 | 116,13 | 164,45 |
| brain lower grade glioma | **LGG** | 139,37 | 135,48 | 153,7 | 99,21 | 108,19 | 110,12 |
| liver hepatocellular carcinoma | **LIHC** | 145,4 | 75,58 | 74,78 | 36,99 | 53,23 | 35,44 |
| lung adenocarcinoma | **LUAD** | 128,07 | 83,36 | 122,2 | 120,82 | 81,34 | 70,18 |
| lung squamous cell carcinoma | **LUSC** | 145,69 | 83,28 | 212,53 | 122,68 | 113,26 | 70,26 |
| ovarian serous cystadenocarcinoma | **OV** | 106,79 | 53,68 | 201,89 | 126,74 | 86,54 | 60,85 |
| pancreatic adenocarcinoma | **PAAD** | 177,9 | 34,75 | 126,01 | 64,83 | 69,37 | 30,51 |
| pheochromocytoma & paraganglioma | **PCPG** | 148,29 | 151,7 | 154,71 | 137,75 | 103,18 | 68,55 |
| prostate adenocarcinoma | **PRAD** | 134,81 | 89,58 | 120,68 | 104,93 | 65,91 | 67,05 |
| rectum adenocarcinoma | **READ** | 300,34 | 82,06 | 181,42 | 170,59 | 112,68 | 102,48 |
| sarcoma | **SARC** | 121,33 | 127,52 | 120,12 | 172,45 | 107,06 | 116,16 |
| skin cutaneous melanoma | **SKCM** | 208,82 | 100,15 | 135,42 | 153,72 | 85,39 | 55,1 |
| stomach adenocarcinoma | **STAD** | 185,45 | 90,07 | 142,22 | 105,86 | 71,46 | 51,95 |
| testicular germ cell tumor | **TGCT** | 72,84 | 48,33 | 142,89 | 118,83 | 126,04 | 125,46 |
| thyroid carcinoma | **THCA** | 99,47 | 76,87 | 118,08 | 109,17 | 63,23 | 98,72 |
| thymoma | **THYM** | 120,15 | 11,24 | 136,55 | 21,67 | 111,2 | 16,29 |
| uterine carcinosarcoma | **UCEC** | 145,39 | 58,84 | 142,03 | 135,35 | 115,94 | 87,44 |
| uterine corpus endometrioid carcinoma | **UCS** | 170 | 55,98 | 170,5 | 134,94 | 115,72 | 87,34 |

**Suppl. Table 2. Expression profiles of VDAC genes from GEPIA server across 31 tumor and paired normal tissues.** No data was found for *VDAC1P5, VDAC1P7, VDAC1P10, VDAC1P12* and *VDAC2P2* pseudogenes. (a) *VDAC1* pseudogenes; (b) *VDAC2* pseudogene; (c) *VDAC3* pseudogene. The median expression of Log2TPM+1 of certain tumor type or normal tissue based on TCGA and GTEX respectively. TPM= transcript per million.

**color gradient**

**Lowest value** — **Highest value**

**(a) *VDAC1* pseudogenes**

| Tumor | Acronym | *VDAC1P1* Tumor | *VDAC1P1* Normal | *VDAC1P2* Tumor | *VDAC1P2* Normal | *VDAC1P3* Tumor | *VDAC1P3* Normal |
|---|---|---|---|---|---|---|---|
| adrenocortical cancer | **ACC** | 0,35 | 0,35 | 0,33 | 0,04 | 0 | 0 |
| bladder urothelial carcinoma | **BLCA** | 0,41 | 0,28 | 0,21 | 0,1 | 0 | 0 |
| breast invasive carcinoma | **BRCA** | 0,64 | 0,29 | 0,15 | 0,04 | 0 | 0 |
| cervical & endocervical cancer | **CESC** | 0,36 | 0,21 | 0,19 | 0 | 0 | 0 |
| cholangiocarcinoma | **CHOL** | 0,46 | 0,12 | 0,25 | 0,07 | 0 | 0 |
| colon adenocarcinoma | **COAD** | 0,75 | 0,28 | 0,47 | 0 | 0 | 0 |
| diffuse large B-cell lymphoma | **DLBC** | 0,33 | 0,03 | 0,18 | 0 | 0 | 0 |
| esophageal carcinoma | **ESCA** | 0,54 | 0,39 | 0,08 | 0,04 | 0 | 0 |
| glioblastoma multiforme | **GBM** | 0,99 | 0,39 | 0,12 | 0 | 0 | 0 |
| head & neck squamous cell carcinoma | **HNSC** | 0,42 | 0,5 | 0,25 | 0,26 | 0 | 0 |
| kidney chromophobe | **KICH** | 0,76 | 0,4 | 0,44 | 0,15 | 0 | 0 |
| kidney clear cell carcinoma | **KIRC** | 0,91 | 0,77 | 0,19 | 0,11 | 0 | 0 |
| kidney papillary cell carcinoma | **KIRP** | 0,47 | 0,41 | 0,22 | 0,14 | 0 | 0 |
| acute myeloid leukemia | **LAML** | 0,23 | 0,78 | 0 | 0,11 | 0 | 0 |
| brain lower grade glioma | **LGG** | 0,37 | 0,39 | 0,22 | 0 | 0 | 0 |
| liver hepatocellular carcinoma | **LIHC** | 0,33 | 0,18 | 0,16 | 0,01 | 0 | 0 |
| lung adenocarcinoma | **LUAD** | 0,38 | 0,3 | 0,2 | 0,04 | 0 | 0 |
| lung squamous cell carcinoma | **LUSC** | 0,49 | 0,3 | 0,15 | 0 | 0 | 0 |
| ovarian serous cystadenocarcinoma | **OV** | 0,3 | 0,17 | 0,05 | 0 | 0 | 0 |
| pancreatic adenocarcinoma | **PAAD** | 0,48 | 0,1 | 0,21 | 0 | 0 | 0 |
| pheochromocytoma & paraganglioma | **PCPG** | 0,48 | 0,35 | 0,15 | 0,13 | 0 | 0 |
| prostate adenocarcinoma | **PRAD** | 0,35 | 0,26 | 0,16 | 0 | 0 | 0 |
| rectum adenocarcinoma | **READ** | 0,82 | 0,26 | 0,54 | 0 | 0 | 0 |
| sarcoma | **SARC** | 0,25 | 0,25 | 0,16 | 0,19 | 0 | 0,01 |
| skin cutaneous melanoma | **SKCM** | 0,49 | 0,3 | 0,3 | 0 | 0 | 0 |
| stomach adenocarcinoma | **STAD** | 0,56 | 0,28 | 0,11 | 0,02 | 0 | 0 |
| testicular germ cell tumor | **TGCT** | 0,2 | 0,14 | 0,06 | 0 | 0 | 0 |
| thyroid carcinoma | **THCA** | 0,35 | 0,33 | 0,12 | 0 | 0 | 0 |
| thymoma | **THYM** | 0,27 | 0,03 | 0,13 | 0 | 0 | 0 |
| uterine carcinosarcoma | **UCEC** | 0,37 | 0,17 | 0,17 | 0 | 0 | 0 |
| uterine corpus endometrioid carcinoma | **UCS** | 0,31 | 0,18 | 0,23 | 0 | 0 | 0 |

**Suppl. Table 3. Genes at the 6q24.2 locus controlled by the same elements in the *VDAC1P8* pseudogene.** Data shown below are from GeneCards database. *From GWAS catalog; **From GEPIA database; ND= no data available; lncRNA=long non-coding RNA; rRNA=ribosomal RNA.

| Gene | Location | Entrez ID | N° Splice variants | Ensembl Biotype | AML phenotype associated |
|---|---|---|---|---|---|
| *ADAT2* | chr6:143,422,832-143,450,695 | 134637 | 3 | 2 alternative coding sequences, 1 processed transcript | YES* |
| *PEX3* | chr6: 143,450,805-143,490,616 | 8504 | 3 | 2 alternative coding sequences, 1 retained intron | NO |
| *RNA5SP221* | chr6: 143,449,809-143,449,911 | 106480759 | 1 | rRNA pseudogene | NO |
| *LTV1* | chr6: 143,843,338-143,863,812 | 84946 | 1 | coding sequence | YES** |
| *PLAGL1* | chr6: 143,940,300-144,064,599 | 5325 | 27 | 21 alternative coding sequences, 6 processed transcript | NO |
| *TUBB8P2* | chr6: 143,436,216-143,436,710 | 106478982 | 1 | pseudogene | YES** |
| *FUCA2* | chr6: 143,494,812-143,511,720 | 2519 | 3 | 2 coding sequence, 1 processed transcript | YES* |
| *ENSG00000270890* | chr6:143,858,062-143,858,689 | ENSG00000270890 | 1 | pseudogene | YES** |
| *ENSG00000278206* | chr6:143,484,979-143,507,327 | ENSG00000278206 | 1 | lncRNA | YES* |
| *CM03496-315* | *chr6:143,484,979-143,507,327* | *N/D* | *N/D* | *ncRNA* | *YES**** |
| *RF00001-290* | chr6:143,449,809-143,449,911 | GC06P143451 | 1 | rRNA | ND |
| *HSALNG0054032* | chr6:143,443,163-143,450,676 | GC06M143443 | 1 | lncRNA | ND |
| *MN298114-199* | chr6:143,103,524-143,530,080 | GC06M143104 | 1 | lncRNA | YES* |

**From GWAS catalog; **From GEPIA database; ***From ENA (European Nucleotide Archive at EBI); ND= no data available; lncRNA=long non-coding RNA; ncRNA= non-coding RNA; rRNA=ribosomal RNA.*

**Suppl. Table 4. BLAST analysis of human *VDAC1P8* pseudogene in all primate species.**

| ORTHOLOGOUS SEQUENCES OF HUMAN *VDAC1P8-201* IN PRIMATES | | | | | | | |
|---|---|---|---|---|---|---|---|
| **Primate** | **Species** | **Coordinates** | **Score** | **ID %** | **E value** | **Coverage %** | **Syntenic** |
| New Word Apes | *Callithrix jacchus* (Marmoset) | 4:146357145-146357996 (+) | 953 | 89 | 0.0 | 99 | Yes |
| Old Word Apes | *Macaca mulata* (Macaque) | 4:47268504-47269350 (+) | 1251 | 94 | 0.0 | 100 | Yes |
| | *Crab-eating macaque* (Macaque) | 4:48083878-48084724 (+) | 1291 | 94 | 0.0 | 100 | Yes |
| | *Mandrillus leucophaeus (Drill)* | KN977571:1371225-1372071 (-) | 1299 | 94 | 0.0 | 100 | Yes |
| | *Olive baboon* (Baboon) | 6:119151340-119152186 (-) | 1275 | 94 | 0.0 | 100 | Yes |
| Anthropomorphic Apes | *Nomascus leucogenys* (Gibbon) | 3:130879543-130880389 (+) | 1397 | 96 | 0.0 | 100 | Yes |
| | *Sumatran orangutan* (Pongo abelii) | 6:142016901-142017751 (+) | 1490 | 97 | 0.0 | 100 | Yes |
| | *Gorilla gorilla* (Gorilla) | 6:146164781-146165636 (+) | 1567 | 98 | 0.0 | 100 | Yes |
| Hominids | *Pan paniscus* (Bonobo) | 6:145987699-145988551 (+) | 1615 | 99 | 0.0 | 100 | Yes |
| | *Pan troglodytes* (Chimpanzee) | 6:147509704-147510556 (+) | 1623 | 99 | 0.0 | 100 | Yes |

**Suppl. Table 5. List of primers used for Real-Time amplification.**

| **Primer** | **Sequence** | |
|---|---|---|
| ***VDAC1*** | FW | 5'-TCCAGCCTGATAGGTTTAGG-3' |
| | REV | 5'-TTCTGAAGGTAGCTATGCTGC-3' |
| ***VDCA1P8-201*** | FW | 5'-TCAGTCCGAGGCACTCTTAAG-3' |
| | REV | 5'-TCACCTTCTGCTAATTGGAGCT-3' |
| ***GAPDH*** | FW | 5'-GAAGGTGAAGGTCGGAGTC-3' |
| | REV | 5'-GAAGATGGTGATGGGATTTC-3' |
